# scQuery: a web server for comparative analysis of single-cell RNA-seq data

**DOI:** 10.1101/323238

**Authors:** Amir Alavi, Matthew Ruffalo, Aiyappa Parvangada, Zhilin Huang, Ziv Bar-Joseph

**Affiliations:** Computational Biology Department, School of Computer Science, Carnegie Mellon University, Pittsburgh, PA 15213; Machine Learning Department, School of Computer Science, Carnegie Mellon University, Pittsburgh, PA 15213

## Abstract

Single cell RNA-Seq (scRNA-seq) studies often profile upward of thousands of cells in heterogeneous environments. Current methods for characterizing cells perform unsupervised analysis followed by assignment using a small set of known marker genes. Such approaches are limited to a few, well characterized cell types. To enable large scale supervised characterization we developed an automated pipeline to download, process, and annotate publicly available scRNA-seq datasets. We extended supervised neural networks to obtain efficient and accurate representations for scRNA-seq data. We applied our pipeline to analyze data from over 500 different studies with over 300 unique cell types and show that supervised methods greatly outperform unsupervised methods for cell type identification. A case study of neural degeneration data highlights the ability of these methods to identify differences between cell type distributions in healthy and diseased mice. We implemented a web server that compares new datasets to collected data employing fast matching methods in order to determine cell types, key genes, similar prior studies, and more.

## Introduction

Single-cell RNA sequencing (scRNA-seq) has recently emerged as a major advancement in the field of transcriptomics (Kolodziejczyk et al., 2015). Compared to bulk (many cells at a time) RNA-seq, scRNA-seq can achieve a higher degree of resolution, revealing many properties of subpopulations in heterogeneous groups of cells (Wills et al., 2013). Several different cell types have now been profiled using scRNA-seq leading to the characterization of sub-types, identification of new marker genes, and analysis of cell fate and development (Lescroart et al., 2018; Patel et al., 2014; Zeisel et al., 2015).

While most work attempted to characterize expression profiles for specific (known) cell types, more recent work has attempted to use this technology to compare differences between different states (for example, disease vs. healthy cell distributions) or time (for example, sets of cells in different developmental time points or age) (Mathys et al., 2017; Rizvi et al., 2017). For such studies, the main focus is on the characterization of the different cell types within each population being compared, and the analysis of the differences in such types. To date, such work primarily relied on known markers (Usoskin et al., 2015) or unsupervised (dimensionality reduction or clustering) methods (Jaitin et al., 2014). Markers, while useful, are limited and are not available for several cell types. Unsupervised methods are useful to overcome this, and may allow users to observe large differences in expression profiles, but as we and others have shown, they are harder to interpret and often less accurate than supervised methods (Lin et al., 2017).

To address these problems, we have developed a framework that combines the idea of markers for cell types with the scale obtained from global analysis of all available scRNA-seq data. We developed scQuery, a web server that supports the analysis of new, large scale scRNA-seq datasets. scQuery relies on scRNA-seq data collected from over 500 different experiments. We developed a common pipeline to uniformly process all scRNA-seq experiments in public databases. The pipeline automatically associates different profiles with cell type based on a constrained ontology and then aligns the raw read data, assigns them to a pre-defined set of genes, and quantifies their expression. Next, we used neural networks to reduce the dimensions of the input data in a supervised way to improve retrieval run time, reduce storage space, and improve accuracy. All profiles are then stored in a web server and newly uploaded data (which is also processed and reduced using the same pipeline) is compared to all database data using fast approximate nearest neighbors methods. The web server then provides users with information about the cell type predicted for each cell, overall cell type distribution, set of differentially expressed (DE) genes identified for cells, prior data that is closest to the new data, and more.

We tested the pipeline, dimensionality reduction, approximate nearest neighbors, and overall web server in several cross-validation experiments. We also performed a case study in which we analyzed close to 2000 cells from a neurodegeneration study (Mathys et al., 2017). As we show, in all cases we observe good performance of the methods we used and of the overall web server for the analysis of new scRNA-seq data.

## Results

We developed a pipeline (Figure 1) for querying, downloading, aligning and quantifying scRNA-seq data. Following queries to the major repositories (Methods) we uniformly processed all datasets so that each was represented by the same set of genes and undergoes the same normalization procedure (RPKM). We next attempt to assign each cell to a common ontology term using text analysis. This uniform processing allowed us to generate a combined dataset that represented expression experiments from more than 500 different scRNA-seq studies, representing 300 unique cell types, and totaling over 60K expression profiles that passed our stringent filtering criteria for both expression quality and ontology assignment (Methods). We next used supervised neural network (NN) models to learn reduced dimension representations for each of the input profiles. We tested several different types of NNs including architectures that utilize prior biological knowledge to reduce overfitting as well as architectures that directly learn a discriminatory reduced dimension profile (Siamese architectures (Koch, Zemel, and Salakhutdinov, 2015)). Reduced dimension profiles for all data were then stored on a web server that allows users to perform queries to compare new scRNA-seq experiments to all data collected so far to determine cell types, identify similar experiments and focus on key genes.

**Figure 1:**
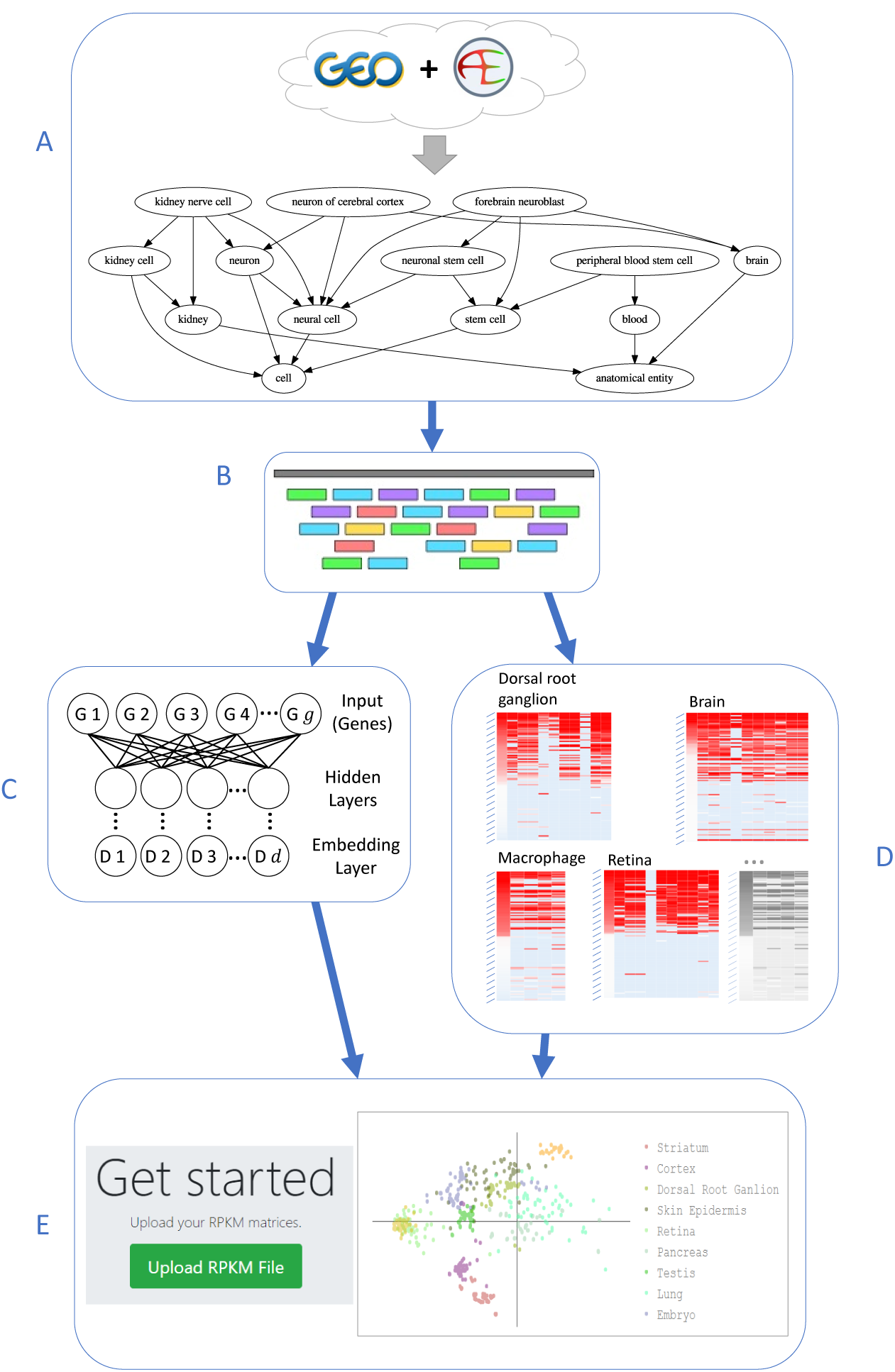
Pipeline for large-scale, automated analysis of scRNA-seq data. **(A)** Bi-weekly querying of GEO and ArrayExpress to download the latest data, followed by automatic label inference by mapping to the Cell Ontology. **(B)** Uniform alignment of all data sets using HISAT2, followed by quantification to obtain RPKM values. **(C)** Supervised dimensionality reduction using our neural embedding models. **(D)** Identification of cell type-specific gene lists using differential expression analysis. **(E)** Integration of data and methods into a publicly available web application.

### Statistics for data processing and downloads

To retrieve all available scRNA-seq data we queried the two largest databases, GEO and ArrayExpress, for scRNA-seq data. Supporting Figure S1 presents screenshots of queries to the NCBI GEO and ArrayExpress databases similar to the ones we used here, though our queries utilized automated APIs instead of the web interfaces shown in these figures. Figure 2 and Supporting Figure S2 respectively show study and cell counts by month, with respect to the “release date” data provided by GEO and ArrayExpress. As can be seen, while cell counts increase over time, there is a lag in availability of raw data and author-processed supplementary data available through NCBI GEO and ArrayExpress systems. Since our pipeline is automated, we expect to be able to collect and analyze much more data over the next several months.

**Figure 2:**
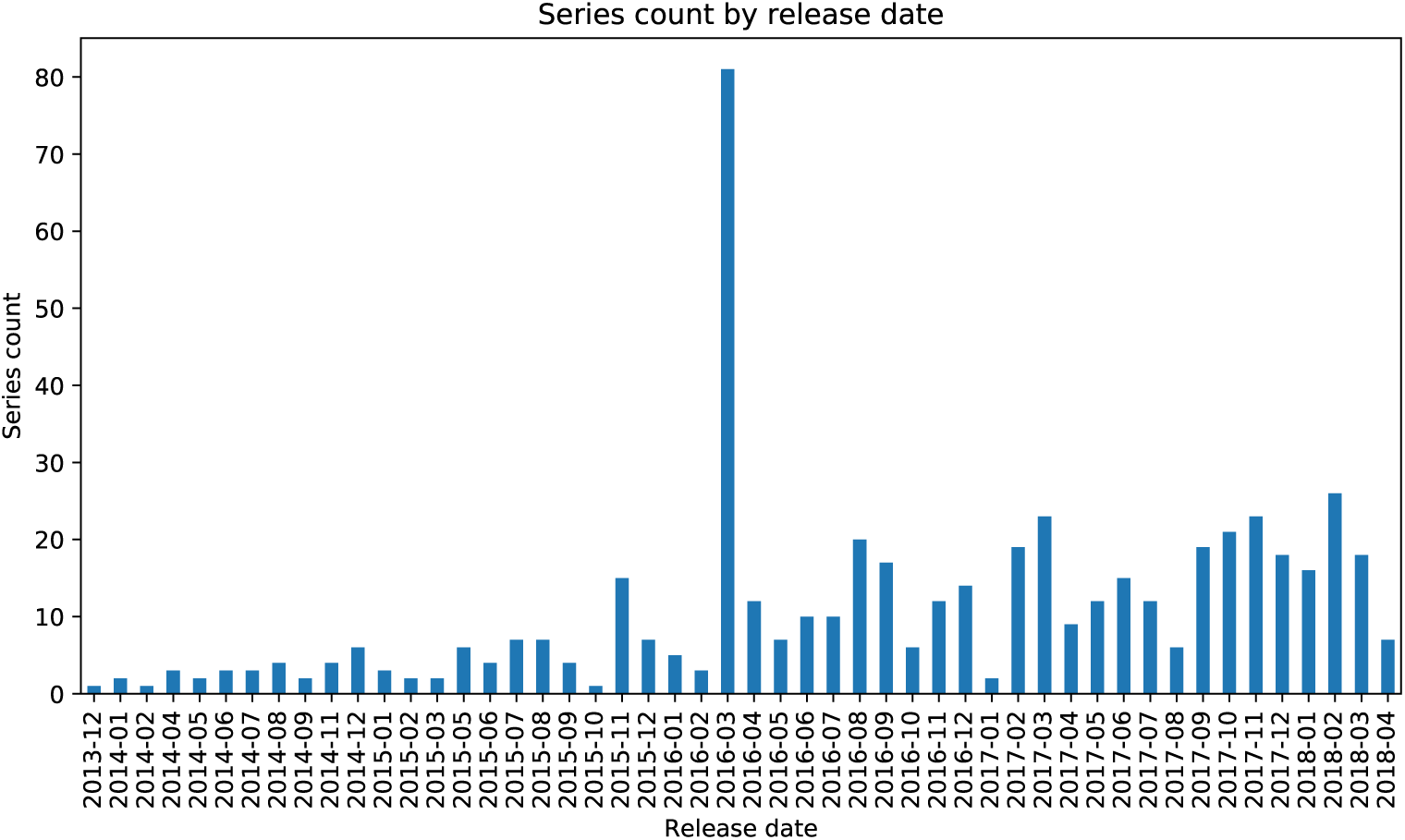
Number of studies released each month in GO and ArrayExpress.

Our “mouse single-cell RNA-seq” query matched a total of 155,242 cells, of which 71,982 have raw data and 29,216 had only author-processed data. We used established ontologies to determine the cell type that was profiled (Methods) for each cell expression dataset we downloaded. Of the 2,356 unique descriptors we obtained for all cells, 1,860 map to at least one term in the cell ontology. Of the 5,010 distinct cell ontology terms (restricted to the CL and UBERON namespaces), 307 are assigned at least one cell expression profile.

Of the 71,982 cells for which raw data is available, 49,237 had alignment rates above our cutoff of 40%. Of these 49,237, we identified 2,473 raw data files that contained reads from multiple cells, but lacked any metadata that allowed us to assign reads to individual cells. This leaves 46,764 cells that are usable for building our scRNA-seq database.

### Neural networks for supervised dimensionality reduction

We trained several different types of supervised and unsupervised (autoencoder) NNs. These included models with the label matching a cell type as the output (with the layer before last serving as the reduced dimension) (Lin et al., 2017), models that directly optimize a discriminatory reduced dimension layer (using as input pairs or triplets of matched and unmatched profiles), and autoencoders which attempt to reconstruct the input by introducing a lower dimension bottleneck. See Supporting Methods Figures S9, S11, and S12 for details. Some of the models utilized prior biological knowledge as part of the architecture to reduce overfitting (including protein-protein and protien-DNA interaction data and hierarchical GO assignments) while others did not (dense, autoencoders). We experimented with various hyperparameters (Methods), and all of our neural network models were trained for up to 100 epochs, with training terminated early if the model converged. All models converged sooner than the full 100 epochs (Supporting Figure S3). Performance on a held out validation set was assessed after each epoch during training. The final weights chosen for each model were those at the end of the epoch with the lowest validation loss out of the 100 epochs. For triplet networks, we also monitored the fraction of “active triplets” in each batch as a selection criterion. Most models trained in minutes (Figure S4), but more complex models, such as our hierarchical GO architectures (116,271,314 parameters) took hours to train (1 hour and 57 minutes for the triplet version).

After training each of our neural embedding models, we evaluated their performance using retrieval testing as described in Methods. Cells used for testing are completely disjoint from the set of cells that were used for training and come from different studies so that batch effects and other experimental artifacts do not affect performance and evaluation. The results of this retrieval testing for a selection of architecture types and cell types are shown in Figure 3. These models have been trained using the data that we processed ourselves in addition to data from studies that only had author-processed data available (these required missing-data imputation, see Methods) though similar results were obtained on models trained using only our own processed data (Supporting Figure S5). It is common to assess retrieval performance with mean average precision (MAP). Here, each value in the table represents the mean average “flexible” precision (MAFP, Supporting Methods), which allows for scores between 0 and 1 for matching a cell type to a similar or parent type (for example, a cortex cell matched to brain, Methods).

**Figure 3:**
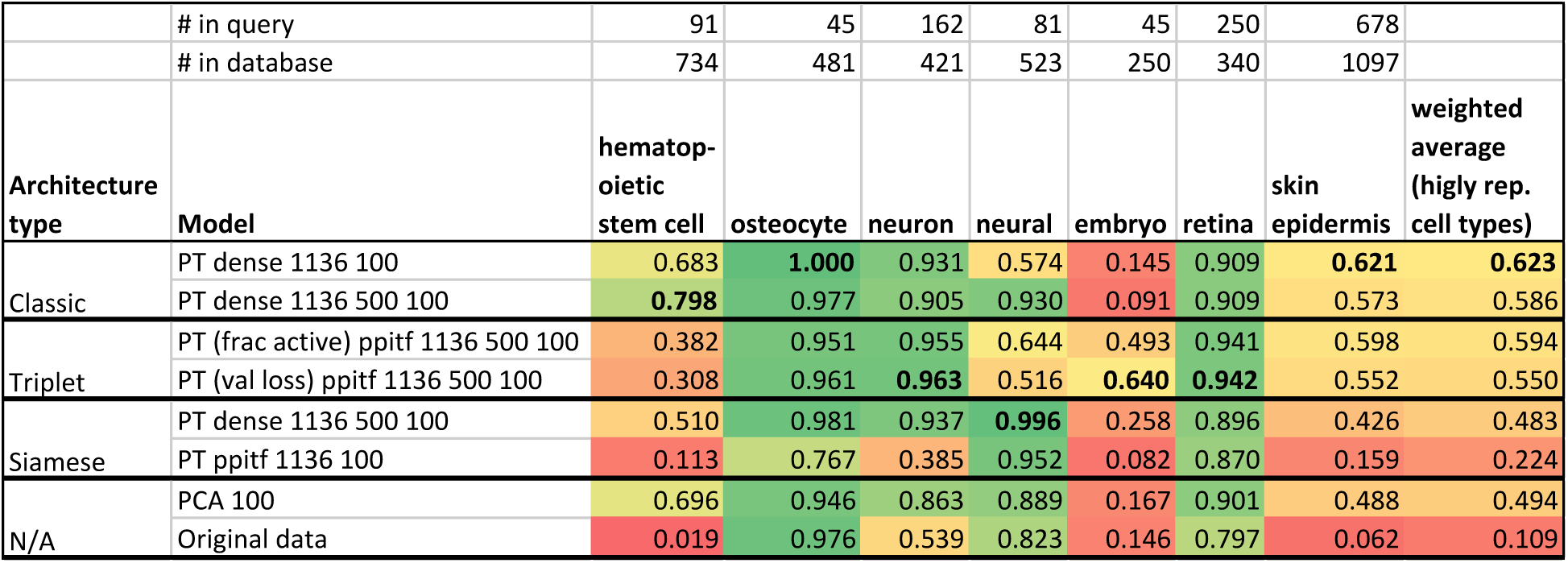
Retrieval testing results of various architectures, as well as PCA and the original (unreduced) expression data. Scores are MAFP values. “PT” indicates that the model had been pretrained using the unsupervised strategy (Supporting Methods). Numbers after the model name indicate the hidden layer sizes. For example, “dense 1136 500 100” is an architecture with three hidden layers. The metrics in parenthesis for the triplet architectures indicate the metric used to select the best weights over the training epochs. For example, “frac active” indicates that the weights chosen for that model were the ones that had the lowest fraction of active triplets in each mini-batch. The final column is the weighted average score over those cell types with at least 1000 cells in the database (some not shown in table), and the weights are the number of such cells in the query set.

The best scoring model achieved a weighted average (accross all cell types) MAFP of 0.576 which is very high when considering the fact that this was a 45-way classification problem (while our database contains over 300 cell types, only 45 types had independent data from multiple studies and these were used for the analysis discussed here). When restricting the analysis to the six cell types for which we had more than 1000 cells in our database, results for this model further improved to 0.623. This top performing model (first row in Figure 3) employed a dense architecture (two-hidden-layer perceptron network). The next best model overall (based on the weighted average) was a similarly defined and performing dense architecture with three-hidden-layers (not shown), followed by a PPITF architecture with three sparsely-connected hidden layers trained as a triplet network (third row in Figure 3). We also see that for specific cell types, other neural networks perform better. Specifically, triplet networks perform best for neuron, embryo, and retina. Siamese architectures perform the best in the neural cell type. We also note that the best performing models were those that were pretrained with an unsupervised strategy (a full table with result from over 100 models, including those from models without pretraining are available on our web server). Finally, as is clear from the last two rows, supervised neural network embeddings consistently outperform PCA and the original data in the retrieval task.

### Gene set enrichment analysis (GSEA) of cell-type specific DE genes

We further used our ontology assignments to identify cell type-specific genes (Methods). For this we used read counts rather than RPKM following several prior methods that determined that such analysis provides a better list of DE genes (Bullard et al., 2010; Robinson and Oshlack, 2010). The differential expression analysis we conducted is based on multiple studies for each cell type. Our procedure performs DE analysis for each study, for each cell type independently and then combines the results. This method ensures that resulting DE genes are not batch or lab related but rather real DE for the specific cell type. We used this method to identify cell type specific genes for 39 cell types. The number of significant (< 0.05 FDR adjusted p-value) DE genes for each cell type ranged from 170 for embryonic stem cells to 7381 for osteocytes. The full list of DE genes can be found on the supporting web server.

To determine the accuracy of the DE genes and to showcase the effectiveness of the automated processing and ontology assignments we performed GSEA using the Gene Ontology (GO) on the set of DE genes for each cell type. Results for a number of the cell types are presented in Figure 4. As can be seen, even though each of the cell type data we used combined multiple studies from different labs, the categories identified for all of the cell types are highly specific indicating that the automated cell type assignment and processing were able to correctly group related experiments.

**Figure 4:**
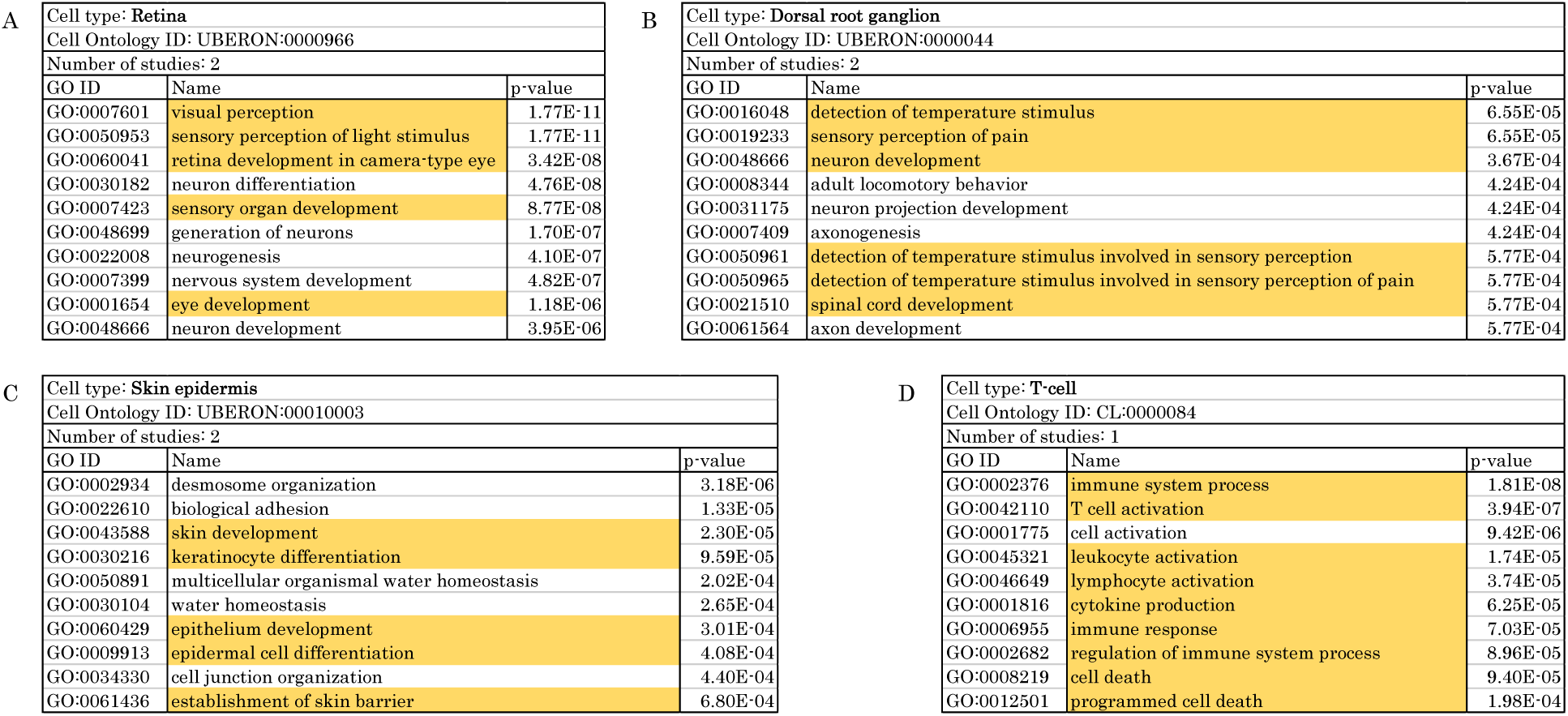
Results of GO enrichment analysis. We used GO: Biological Process as the source gene set, and using the top 50 differentially expressed genes from our cell type-specific gene lists for **(A)** Retina, **(B)** Dorsal root ganglion, **(C)** Skin epidermis, and **(D)** T cell. The top 10 GO terms are shown for each cell type, sorted by FDR adjusted p-values.

As an example, the top three enriched terms for “retina” are “visual perception” (*p* = 1.77 × 10^−11^),“sensory perception of light stimulus” (*p* = 1.77 × 10^−11^), and “retina development in camera-type eye” (*p* = 3.42 × 10^−8^). Cells of dorsal root ganglion are sensory neurons as reflected in Figure 4B with terms such as “detection of temperature stimulus” (*p* = 6.55 × 10^−5^) and “sensory perception of pain” (*p* = 6.55 × 10^−5^). In Figure 4D, nine of the top ten terms are related to immune response and specific aspects of the T cell-mediated immune system. Complete results are available from the supporting web server.

### Query and retrieval

To enable users to compare new scRNA-seq data to the data we processed, and to determine the composition of cell types in such samples we developed a web application. After processing their data (Methods) users can uploaded it to the server. Next, uploaded data is compared to all studies stored in the database. For this, we use approximate nearest neighbor approaches to match these to the data we have pre-processed. Embedding in the reduced dimension representation and the fast matching queries of thousands of cells against hundreds of thousands of database cells can be performed in minutes.

Figure 5 present an example of a partial analysis of newly uploaded data by the web server. The web server clusters cells based on their matched types (Figure 5A), plots their 2D embedding with respect to all other cell types in the database (Figure 5B), highlights the top represented ontology terms in the uploaded cells (Figure 5C), and provides additional information about key DE (Figure 5D) genes and specific studies that it matched to the uploaded data (Figure 5E).

**Figure 5:**
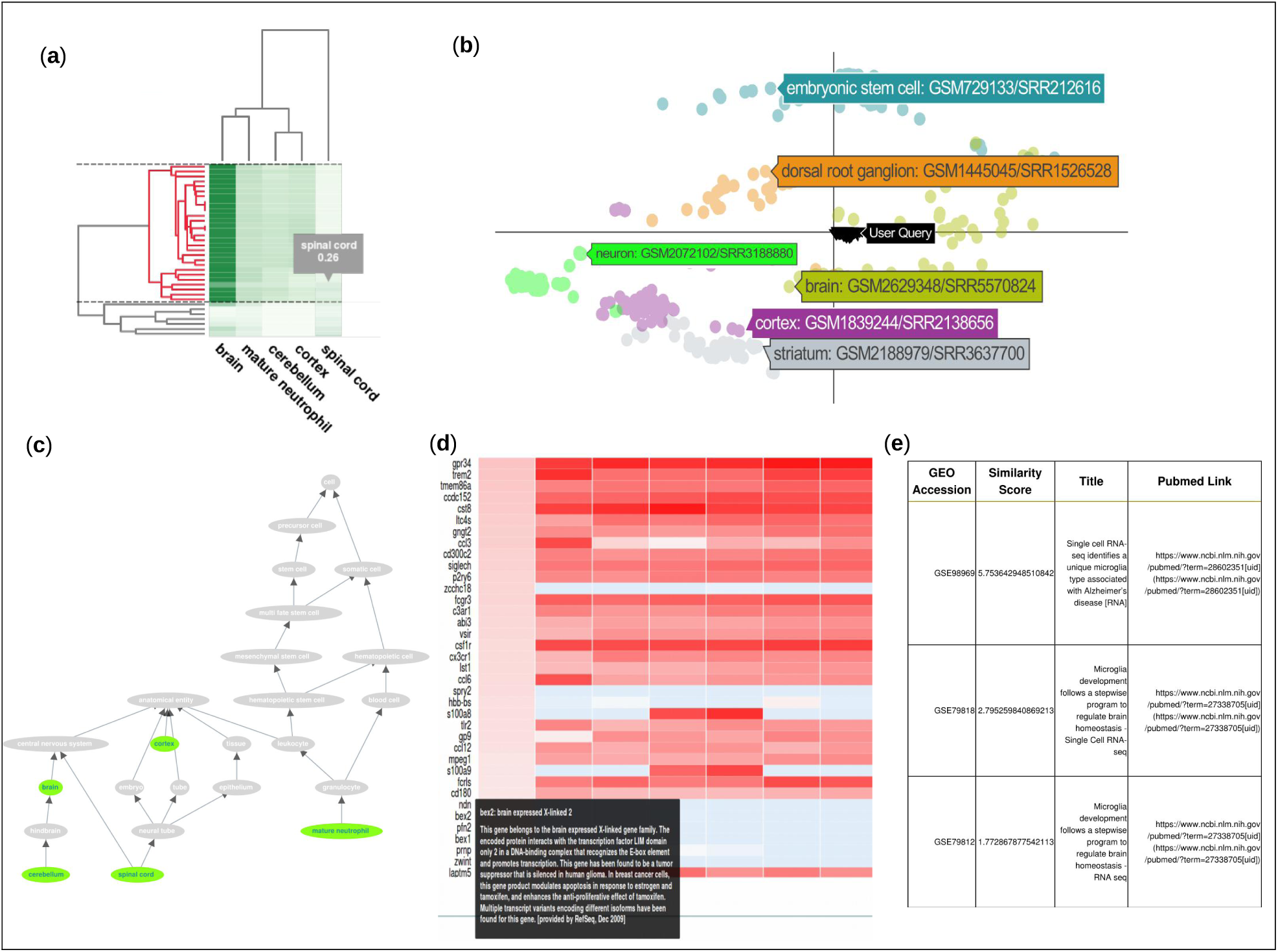
The scQuery web server. **(A)** Cluster heatmap of the nearest neighbor results for a query consisting of 40 “brain” and 10 “spinal cord” cells. The horizontal dashed lines demarcate the currently selected cluster and the corresponding dendrogram sub-cluster is highlighted in red. **(B)** 2D scatter plot of the selected sub-cluster (shown as inverted triangles and tagged as “User Query”) along with a handful of other cell-types whose tags show cell-type information and GEO submission ids for a single cell from each cluster. **(C)** Ontology DAG depicting the retrieved cell-types in green while the nodes in gray visualize the path to the root nodes (which reflects paths of cellular differentiation as well as other biological relationships). **(D)** Heatmap displaying log-transformed RPKM values for differentially expressed genes of a subset of the submitted query cells. The first column shows log-tranformed RPKM counts of the same genes averaged over all the cells in the processed database that are mapped to the ontology term activated in the ontology DAG(“brain” here). **(E)** Metadata table for the retrieved hits displaying the GEO accession id, similarity score, publication titles, and their respective pubmed links.

### Mouse brain case study

To test the application of our pipeline we used it to study a recent scRNA-seq neurodegeneration dataset that was not included in our database (Mathys et al., 2017). This study profiled 2208 microglial cells extracted from the hippocampus of the CK-p25 mouse model of severe neurodegeneration. In the CK-p25 mouse model, induction of the p25 gene, a calpain cleaved kinase activator, results in Alzheimer’s disease-like pathology. In the original study, the microglial cells were extracted from control and CK-p25 mice from four time points: before p25 induction (three months old), and one, two, and six weeks after induction (three months 1 week, three months 2 weeks old, and 4 months 2 weeks old, respectively). The goal of the study was to compare the response of microglial cells to determine distinct molecular sub-types, uncover disease-stage specific states, and further characterize the heterogeneity in microglial response. We used the raw read data to perform alignment and quantification (Methods) resulting in 1990 cells that passed our alignment thresholds and were used for the analysis that follows.

### Neural embeddings

We used this data to test several aspects of the method, pipeline, and website. The web server was able to perform a complete analysis of the roughly 2000 cells in minutes. We first compared the supervised (using NN) and unsupervised dimensionality reduction. The cells were transformed to a lower dimensional space using the “PT dense 1136 100” NN followed by t-SNE to get them to 2 dimensions. We compared this to a completely unsupervised dimensionality reduction, as was done in the original paper. Supporting Figure S6 presents the results of this analysis. We observe that the supervised method is able to better account for the differences between the two populations of healthy and disease cells.

### Cell type classifications

We next performed retrieval analysis by using the mouse brain cells as queries against our large database of labeled cells. We classified each query cell based on the most common label in its 100 nearest neighbors in the database (Methods). The results of this cell type classification can be seen in Figure 6.

**Figure 6:**
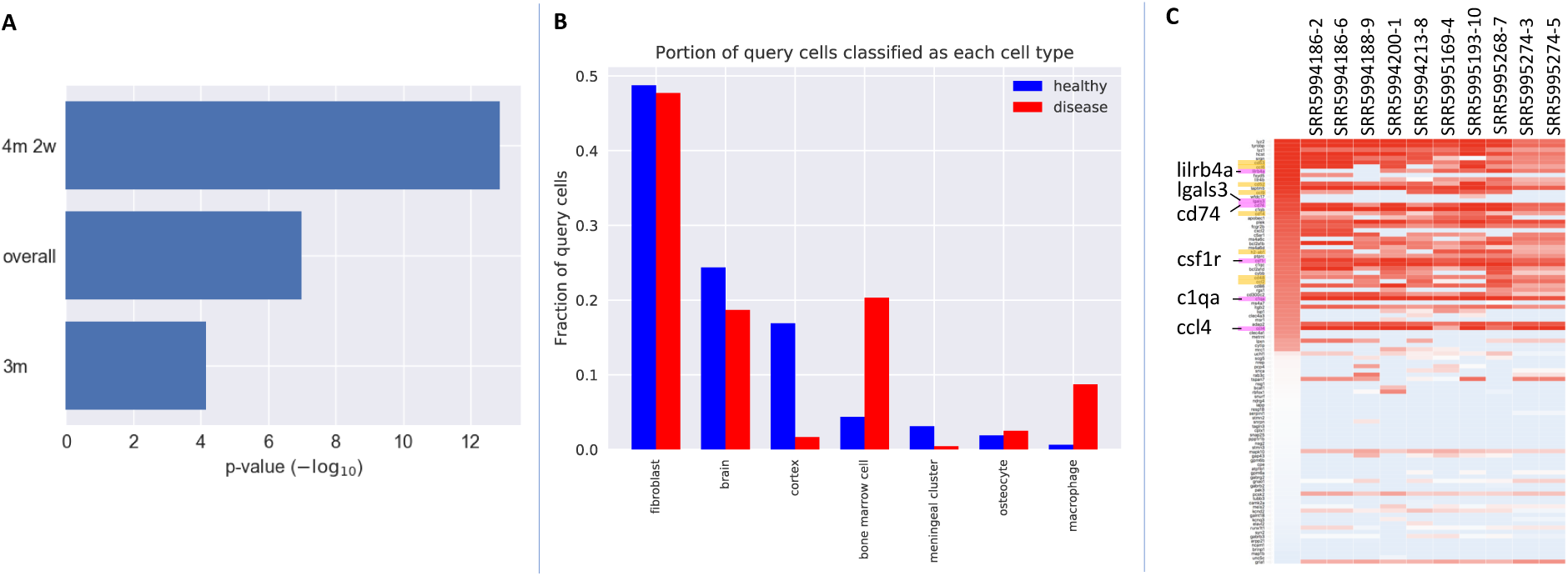
Analysis of mouse neurodegeneration dataset, late response cells. **(A)** p-values of the difference in cell type classification distributions for different time points. Three months was the initial time point in the study, and four months two weeks was the last time point. “Overall” is the pool of all 1990 cells. The p-values are from conducting Fisher’s exact test (for “overall”, the p-value was simulated based on 1e+07 replicates). **(B)** Classification distribution for late stage cells (four months two weeks). **(C)** logRPKM expression of the macrophage-specific genes (Methods) within a cluster of late-stage neurodegenerative cells enriched for “macrophage” labeling by our retrieval server. The leftmost column is the average logRPKM among our database macrophage cells. Genes highlighted in magenta were also found to be up-regulated in the original study (Mathys et al., 2017). Genes highlighted in yellow are additional genes related to immune response that are up-regulated in the query cells. Genes highlighted in yellow in order of top to bottom: Cd53, Ccl6, Cd52, Ccl9, Cd14, H2-ab1, Cd48, Ccl2

We next compared the cell assignments for 3 different groups. An early time point (three months old) in which the healthy and disease mouse models are not expected to diverge, a later time point (four months 2 weeks old) in which differences are expected to be pronounced, and all data collected from the healthy and disease data. As can be seen in Figure 6A, overall assignment indeed reflects these stages with a much more significant difference for the later time point compared to the earlier one, with the entire dataset (which includes more intermediate points) in the middle. Focusing on the later time point, Figure 6B shows the distribution for the latest time point. Several of the cell types identified by the method correspond to brain cells (brain, cortex, meningeal cluster) while others are related to blood and immune response (bone marrow, macrophage). The most common classification among the query cells was “fibroblast.” Recent studies have shown that fibroblast-like cells are common in the brain (Park et al., 2012), and that brain fibroblast cells can express neuronal markers (Vanlandewijck et al., 2018).

As can be seen, the main difference observed between the disease and healthy mice is the increase in the immune system related types of “bone marrow” and “macrophage” cells in the disease model. We believe that while the method labeled these cells as macrophages, they are actually microglia cells which were indeed the cells the authors tried to isolate. To confirm this, we analyzed sets of marker genes that are distinct for macrophages and for microglia (Hickman et al., 2013). Figure S7 shows that indeed, for the cells identified by the method the expressed markers are primarily microglia markers.

The main reason that the method identified them as macrophages is the lack of training data for microglial cells in our database (our train data of high-confidence cell types contains no microglial cells and 273 macrophage cells; our full database contains only 44 microglial cells compared to 603 macrophage cells). Still, the result that disease samples contain more immune cells, which is only based on our analysis of scRNA-seq data (without using any known immune markers) indicates that as more scRNA-seq studies are performed and entered into our database, the accuracy of the results would increase.

### Comparison to our differentially expressed genes for macrophage

We further characterized the gene expression within these microglial cells by comparing the gene expression in the query cells to our cell-type-specific marker genes (Methods). We uploaded the RPKM gene expression values for 256 microglial cells from late-stage neurodegenerative mice (6 weeks after p25 induction) to our web server (Mathys et al., 2017). Given the hits from the retrieval database, we then selected a cluster of these that were enriched for “macrophage”, and then viewed the expression (logRPKM) of our previously calculated cell-type-specific marker genes as a heatmap for a subset of these cells. A screen shot of the interactive heatmap provided by the web server is shown in Figure 6C, where we see that there is rough agreement in up and down regulation trends of the set of macrophage-specific DE genes we identified between the cells in the database (leftmost column) and the query (microglial) cells. We would expect to see this expression pattern for these genes in these cells, as microglia are distinct from macrophages, but are related in function.

Among the up-regulated genes selected for “macrophage” in the web server, we see genes that Mathys et al., 2017 also found to be up-regulated in late-response clusters including a microglial marker (Csf1r), a gene belonging to the chemokine superfamily of proteins (Ccl4), a major histocompatability complex (MHC) class II gene (Cd74), and other genes related to immune response (Lilrb4a, Lgals3), (magenta highlights in Figure 6C). We also see that a different microglial marker (C1qa) (Zhang et al., 2014) is up-regulated. In addition to re-identifying genes of interest from the original study, our method is also able to highlight additional genes that are biologically relevant. These are highlighted in yellow in Figure 6C and include more chemokines (Ccl2, Ccl6, Ccl9), another MHC class II gene (H2-ab1), and other cell surface antigens (Cd14, Cd48, Cd52, Cd53).

## Discussion

We developed a computational pipeline to process all scRNA-seq data deposited in public repositories. We have identified over 500 studies of scRNA-seq data. For each we attempted to download the raw data and to assign each cell to a restricted ontology of cell types. For cells for which this information existed we uniformly processed all reads, ran them through a supervised dimensionality reduction method based on NNs and created a database of cell type profiles. Using the scQuery server, users can upload new data, process it in the same way, and then compare it to all collected scRNA-seq data.

In addition to cell assignments, the web server allows users to view the metadata on which the assignment is based, view the ontology terms that are enriched for their data and the distribution it predicts, and compare the expression of genes in the new profiled cells to genes identified as DE for the various cell types. The web server also clusters the cells and plots a 2D dimensionality reduction plot to compare the expression of the users’ cells with all prior cells types it stores. Applying the method to analyze recent neurodegeneration data led to the identification of significant differences between cell distributions of healthy and diseased mouse models, with the largest observed difference being the set of immune related cells that are more prevalent in the diseased mouse. Our method also revealed additional up-regulated immune-related genes in the late-stage neurodegenerative cells that were classified as “macrophage” cells.

While the pipeline was able to process several of the datasets we identified on public repositories, not all of them could be analyzed. Specifically, many studies lacked raw scRNA-seq reads, and thus could not be processed via our uniform expression quantification pipeline. Though we were able to find author-processed expression data for many studies, usage of this data is complicated by different gene selections, data format differences (*e.g.* RPKM vs. FPKM vs. TPM vs. read counts), and more. Additionally, several studies profiled thousands of cells but published far fewer raw data files, with each raw data file containing reads from hundreds or thousands of cells but no metadata that allows each read to be assigned to a unique cell.

In addition to issues with processing data that has already been profiled and deposited, we observed that cell type distribution in our database is still very skewed. While some cell types are very well represented (“bone marrow cell”: 6,283 cells, “dendritic cell”: 4,126 cells, “embryonic stem cell”: 2,963 cells) others are either completely missing or were only represented with very few samples (“leukocyte”: 12 cells, “B cell”: 22 cells, “microglial cell”: 44 cells, “cardiac muscle cell”: 72 cells). Such skewed distribution can cause challenges to our method leading to cells being assigned to similar, but not the correct, types. These are still the early days of scRNA-seq analysis with several public and private efforts to characterize cell types more comprehensively. Our dataset retrieval and processing pipeline (including cell type assignments) is fully automated and we expect that once more experiments are available they would be added to the database and server. We believe that as more data accumulates the accuracy of scQuery would increase making it the tool of choice for cell type assignment and analysis.

## Acknowledgements

Work partially supported by NIH grant 1R01GM122096 and a James S. McDonnell Foundation Scholars Award in Studying Complex Systems to Z.B.-J. We thank Hongyu Zheng (Computational Biology Dept., CMU) for his work on cell-type similarity calculation. We also thank Andreas Pfenning for his suggestions and advice on the mouse brain case study analysis.

## Author Contributions

Conceptualization, Z.B.-J.; Methodology, Z.B.-J., A.A., and M.R.; Software, A.A., M.R., A.P., and Z.H.; Formal Analysis, A.A.; Investigation, Z.B.-J., A.A., M.R.; Data Curation, M.R. and Z.H.; Writing-Original Draft, Z.B.-J., A.A., M.R., A.P., and Z.H.; Writing-Review & Editing, Z.B.-J., A.A., M.R., A.P., and Z.H.; Visualization: A.A., M.R., and A.P.; Supervision: Z.B.-J., A.A., and M.R.; Project Administration, Z.B.-J.; Funding Acquisition, Z.B.-J.

## Methods

### Data collection and preprocessing

We selected a mouse gene set of interest based on the NCBI Consensus CDS (CCDS) which contains 20,499 distinct genes. For genes with multiple isoforms, we consolidate all available coding regions (Supporting Methods). To search for scRNA-seq datasets, we queried the NCBI Gene Expression Omnibus (GEO) and the ArrayExpress database for mouse single-cell RNA-seq series (See Supporting Methods for the queries we used). We then download metadata for each series returned in this query and parse this metadata to identify the distinct samples that comprise each series. We examined the metadata for each sample (*e.g.* library strategy, library source, data processing) and exclude any samples that do not contain scRNA-seq data.

We next attempted to download each study’s raw RNA-seq reads and for those studies for which this data is available we developed a pipeline that uniformly processed scRNA-seq data. We use the reference mouse genome from the UCSC genome browser (Rosenbloom et al., 2014) (build mm10), and align RNA-seq reads with HISAT2 (Kim, Langmead, and Salzberg, 2015) version 2.1.0. We align reads as single-or paired-end as appropriate, and discard samples for which fewer than 40% of reads align to coding regions. We represent gene expression using RPKM. Code for our alignment/quantification pipeline is available at https://github.com/mruffalo/sc-rna-seq-pipeline.

### Labeling using Cell Ontology terms

We use the Cell Ontology (CL) (Bard, Rhee, and Ashburner, 2005) available from http://obofoundry.org/ontology/cl.html to identify the specific cell types which are represented in our GEO query results. We parsed the ontology terms into a directed acyclic graph structure, adding edges between terms for “is a” and “part of” relationships. Note that this choice of edge direction means that all edges point *toward* the root nodes in the ontology.

We use the name and any available synonyms for each ontology term to automatically identify the matching terms for each sample of interest (Supporting Methods). This produces a set of ontology term hits for each sample. We filter these ontology term hits by excluding any terms that are descendants of any other selected terms (*e.g.*, term CL:0000000 “cell” matches many studies), producing a set of “specific” ontology terms for each sample – for any two nodes *u* and *v* in such a set, neither *u* nor *v* is a descendant of the other in the ontology.

### Dimensionality Reduction

Most current analysis methods for scRNA-seq data use some form of dimensionality reduction to visualize and analyze the data, most notably PCA and similar methods (Pierson and Yau, 2015; Yau, 2016) and t-SNE (Tirosh et al., 2016). Past work has shown that while such methods are useful, supervised methods for dimensionality reduction may improve the ability to accurately represent different cell types (see Lin et al., 2017).

Using neural networks for dimensionality reduction has been shown to work well as a supervised technique to learn compact, discriminative representations of data (Hinton and Salakhutdinov, 2006). The original, unreduced dimensions form the input layer to a neural network, where each dimension is an input unit. After training the model towards a particular objective (such as classification), the last hidden layer, which is typically much smaller in the number of units than the input layer, may be taken as a reduced dimensionality representation of the data. These learned features are referred to as *neural embeddings* in the literature, and here we tested a number of different neural network architectures which either explicitly optimize these neural embeddings (for example, Siamese (Chopra, Hadsell, and LeCun, 2005) and triplet networks (Schroff, Kalenichenko, and Philbin, 2015)) or those that only optimize the label accuracy. All neural networks we used were implemented in Python using the Keras API (Chollet, 2015), and our code is available at: https://github.com/AmirAlavi/scrna_nn.

### Neural network architectures

Prior work showed that sparsely connected NN architectures based on protein interaction data can be more effective in determining cell types when compared to dense networks (Lin et al., 2017). Here we further studied other NN networks architectures and compared their performance to the PPI and dense networks. First, we looked at another method to group genes based on the Gene Ontology (GO) (Consortium, 2017). To construct a hierarchical neural network architecture that mirrors the structure of GO we associate input genes with GO nodes. Multiple genes are associated (and connected to) the same node. We use this grouping of the input genes as the first hidden layer of a neural network. Nodes in the next hidden layer will be constructed from GO nodes that are descendants of nodes in the prior layer. We continue this process until the last hidden layer has the desired number of nodes (the size of our reduced dimension). The final result is the network depicted in Figure S12. See also Supporting Methods.

### Siamese architectures trained with contrastive loss

The NNs discussed above indirectly optimize the neural embedding layer by optimizing a classification target function (correct assignment of scRNA-seq data to cell types). A number of NN architectures have been proposed to explicitly optimize the embedding itself. For example, Siamese neural networks (Chopra, Hadsell, and LeCun, 2005; Koch, Zemel, and Salakhutdinov, 2015) (Figure S9) consist of two identical “twin” subnetworks which share the same weights. The outputs of both subnetworks are connected to a “conjoined” layer (sometimes referred to as the “distance” layer) which directly calculates a distance between the embeddings in the last layers of the twin networks. The input to a Siamese network is a pair of data points and the output which is optimized is whether they are similar (same cell type) or not. The loss is computed on the output of the distance layer, and heavily penalizes large distances between items from the same class, while at the same time penalizing small distances between items from different class. Specifically the network optimizes the following loss function:

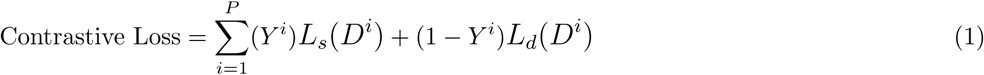

where:

***P*** is the set of all training examples (pairs of data points)

***Y*** is the corresponding label for each pair (1 indicates that the pair belong to the same class, 0 indicates that each sample in the pair come from different classes)

***D*** is the Euclidean distance between the points in the pair computed by the network

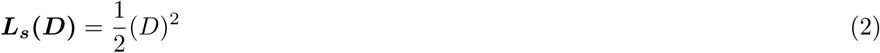

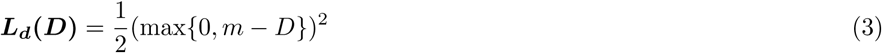

***m*** is a margin hyperparameter, usually set to 1

Following the same motivations as Siamese networks, triplet networks also seek to learn an optimal embedding but do so by looking at three samples at a time instead of just two as in a Siamese network. The triplet loss used by Schroff et al. considers a point (anchor), a second point of the same class as the anchor (positive), and a third point of a different class (negative) (Schroff, Kalenichenko, and Philbin, 2015). See Supporting Methods for details.

### Training and testing of neural embedding models

We conduct supervised training of our neural embedding models using stochastic gradient descent. Although our processed dataset contains many cells, each with a set of labels, we train on a subset of “high confidence” cells to account for any label noise that may have occurred in our automatic term matching process. This is done by only keeping terms that have at least 75 cells mapping to them, and then only keeping cells with a single mapping term. This led to a training set of 21,704 cells from the data we processed ourselves (36,473 cells when combined with author-processed data). We experimented with tanh, sigmoid, and ReLU activations, and found that tanh performed the best. ReLU activation is useful for helping deeper networks converge by preventing the vanishing gradient problem, but here our networks only have a few hidden layers, so the advantage of ReLU is less clear. We also experimented with different learning rates, momentums, and input normalizations (see web server for full results).

Since our goal is to optimize a discriminative embedding, we test the quality of our neural embeddings with retrieval testing, which is similar to the task of cell type inference. In retrieval testing, we query a cell (represented by the neural embedding of its gene expression vector) against a large database of other cells (which are also represented by their embeddings) to find the query’s nearest neighbors in the database.

We separate the **studies** for each cell type when training and testing so that the test set is completely independent of the training set. We find all cell types which come from more than one study, and hold out a complete study for each such cell type to be a part of the test set. Cell types that do not exist in more than one study are all kept in the training set. For our integrated dataset, our training set contained 45 cell types, while our query set was a subset of 26 of the training cell types. After training the model using the training set, the training set can then be used as the database in retrieval testing.

### Evaluation of classification and embeddings for scRNA-seq data

In both training and evaluation of our neural embedding models, we are constantly faced with the question of how similar two cell types are. A rigid (binary) distinction between cell types is not appropriate since “neuron”, “hippocampus”, and “brain” are all related cell types, and a model that groups these cell types together should not be penalized as much as a model that groups completely unrelated cell types together. We have thus extended the NN learning and evaluation methods to incorporate cell type similarity when learning and testing the models. See Supporting Methods for details on how these are used and how they are obtained.

### Differential expression for cell types

We use the automated scRNA-seq annotations we recovered to identify a set of differentially expressed genes for each cell type. Unlike prior methods that often compare two specific scRNA-seq datasets, or use data from a single lab, our integrated approach allows for a much more powerful analysis. Specifically, we can both focus on genes that are present in multiple datasets (and so do not represent specific data generation biases) and those that are unique in the context of the ontology graph (i.e. for two brain related types, find genes that distinguish them rather than just distinguishing brain vs. all others).

Our strategy, presented in Algorithm **??** is DE-method agnostic, meaning that we can utilize any of the various DE tools that exist. In practice we have used SCDE here. This method builds an error model for each cell in the data, where the model is a mixture between a negative binomial and a Poisson (for dropout events) distribution, and then uses these error models to identify differentially expressed genes (Kharchenko, Silberstein, and Scadden, 2014).

Another key aspect of our strategy is the use of meta-analysis of multiple DE experiments. The algorithm attempts to make the best use of the integrated dataset by doing a separate DE experiment for each study that contains cells of a particular cell type, and then combines these results into a final list of DE genes for the cell type. See Supporting Methods for the details of this meta-analysis.

### Large scale query and retrieval

To enable users to compare new scRNA-seq data to the public data we have processed, and to determine the composition of cell types in such samples, we developed a web application. Users download a software package available on the website to process SRA/FASTQ files. The software implements a pipeline that generates RPKM values for the list of genes used in our database and can work on a PC or a server (Supporting Methods).

Once the user processes their data, the data is uploaded to the server and compared to all studies stored in the database. For this, we first use the NN to reduce the dimensions of each of the input datasets and then use approximate nearest neighbor approaches to match these to the data we have pre-processed as we discuss below.

Since the number of unique scRNA-seq expression vectors we store is large, an exact solution obtained by a linear scan of the dataset for the nearest neighbor cell-types would be too slow. To enable efficient searches, we benchmarked three approximate nearest neighbor libraries: NMSLib (Boytsov and Naidan, 2013), ANNoY^1^, and FALCONN (Andoni et al., 2015). Benchmarking revealed that NMSLib was the fastest method (Supporting Figure S8). NMSLib supports optimized implementations for cosine similarity and L2-distance based nearest neighbor retrieval. The indexing involves creation of hierarchical layers of proximity graphs. Hyperparameters for index building and query runtime were tuned to trade-off a high accuracy with reduced retrieval time. For NMSLib, these were: M = 10, efConstruction = 500, efSearch = 100, space =“cosinesimil”, method = “hnsw”, data type = nmslib.DataType.DENSE VECTOR, dtype = nmslib.DistType.FLOAT. Time taken to create the index: 2.6830639410000003 secs. Hyperparameters tuned for the ANNOY library were: number of trees = 50, search k var = 3000. Time taken to create the index: 1.3495307050000065 secs. For FALCONN, a routine to compute and set the hyperparameters at optimal values was used. This calibrates K (number of hash functions) and last cp dimensions. Time taken to create the index: 0.12065599400011706 secs.

### Visualizing query results

We use the approximate nearest neighbors results to compute a similarity measure of each query cell to each ontology term. This is done by identifying the 100 nearest neighbors for each cell and determining the fraction of these matches that belong to a specific cell type. This generates a matrix of similarity measure entries for all query cells against all cell types which is presented as a hierarchical clustering heat-map (Figure 5A). All visualizations are based on this matrix.

We also perform further dimensionality reduction of the query via PCA to obtain a 2D nearest-neighbor style visualization against all cell types in the database and generate the ontology subgraph that matches the input cells. Users can click on any of the nodes in that graph to view the cell associated with it, DE genes related to this cell type, and their expression in the query cells.

In addition to matching cells based on the NN reduced values, we also provide users with the list of experiments in our database that contain cells that are most similar to a subset of uploaded cells the user selects. This provides another layer of analysis beyond the automated (though limited) ontology matching that is based on the cell types extracted for the nearest neighbors.

Finally, users can obtain summary information about cell type distribution in their uploaded cells and can find the set of cells matched to any of the cell types in our database.

## Supporting figures

**Figure S1:**
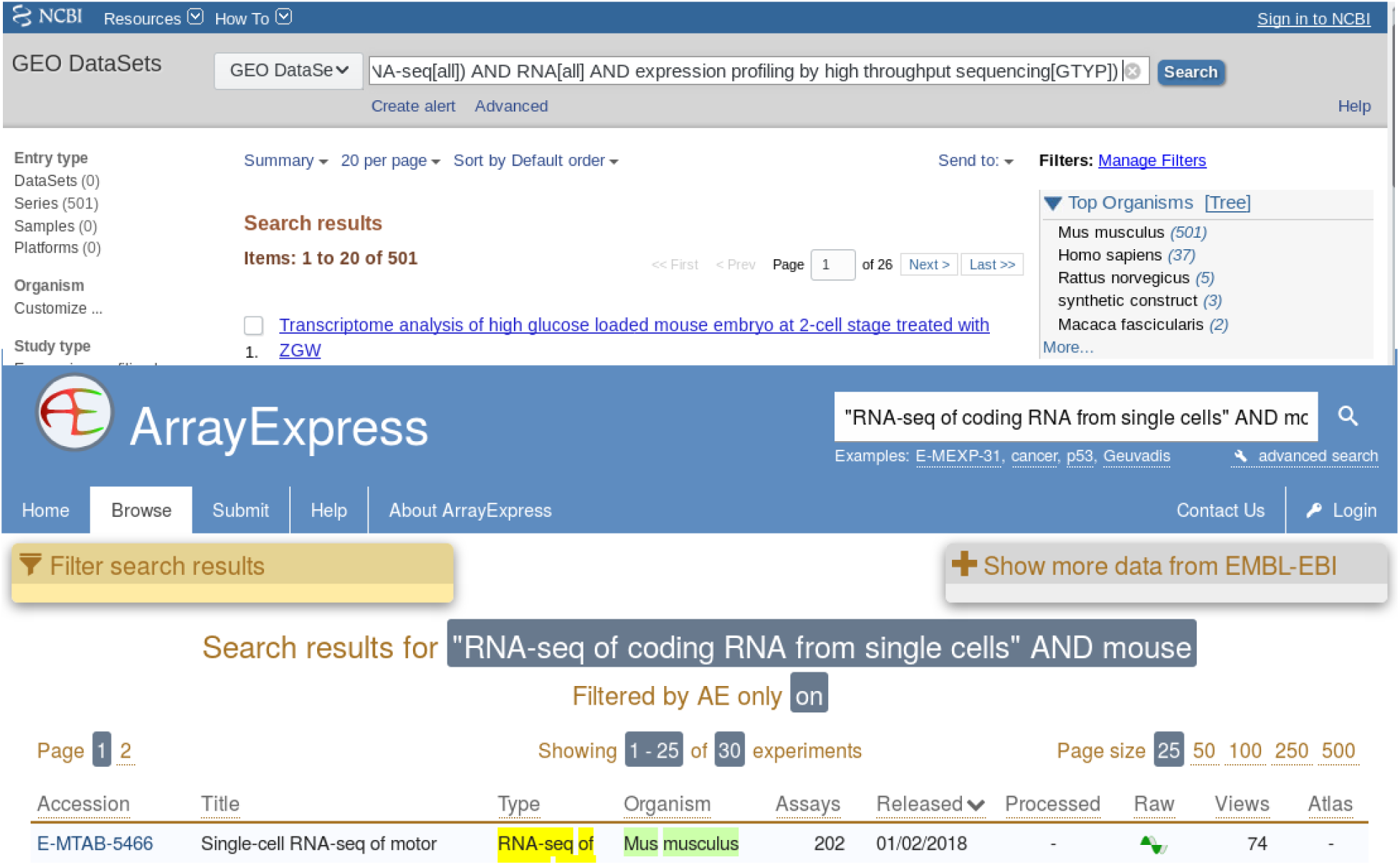
Our queries to the NCBI GEO and ArrayExpress systems, selecting mouse single-cell RNA-seq data.

**Figure S2:**
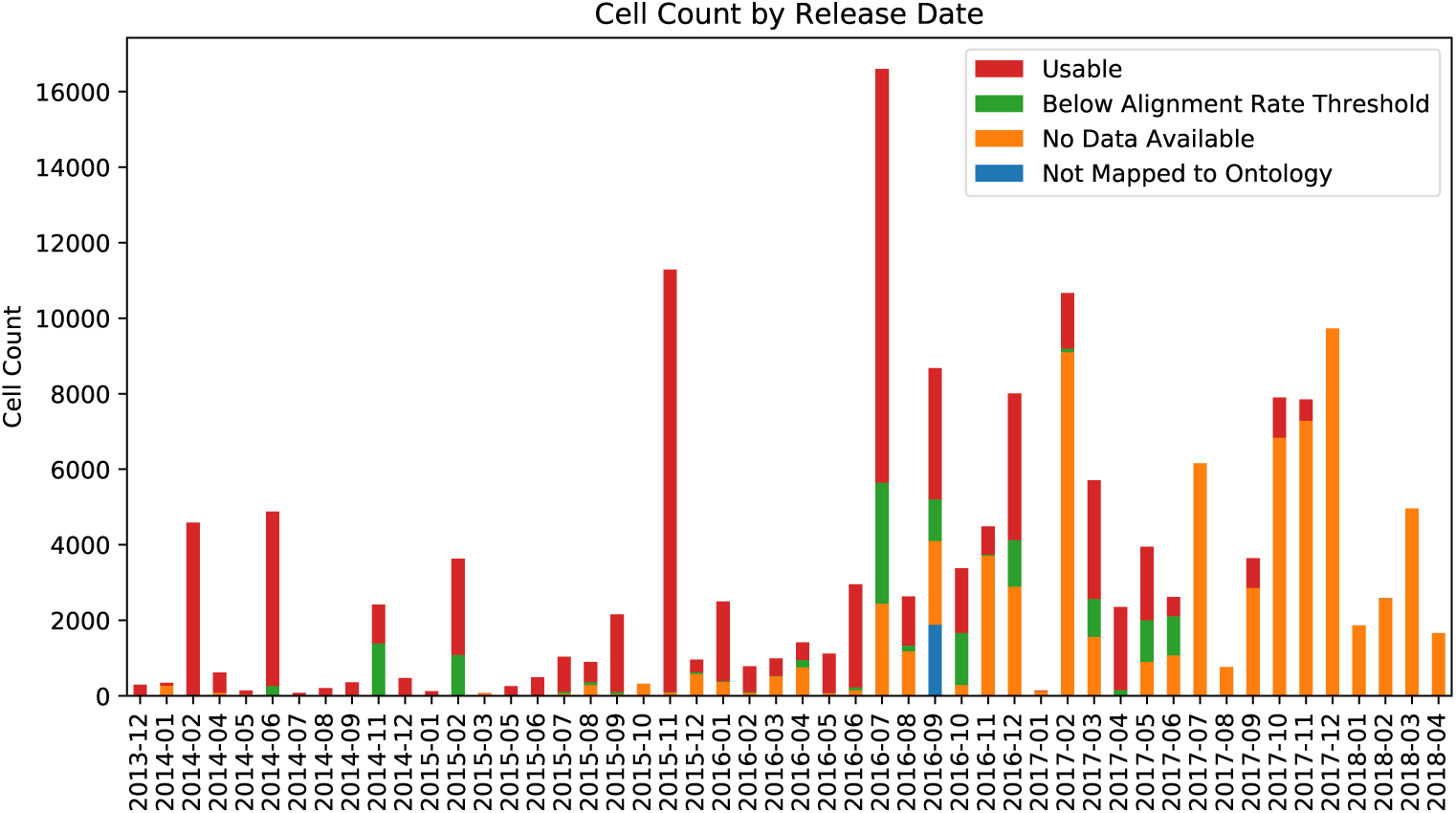
Cell counts by month, separated into four categories: usable, below our alignment rate threshold, no raw or author-processed data available, and unmapped to ontology terms.

**Figure S3:**
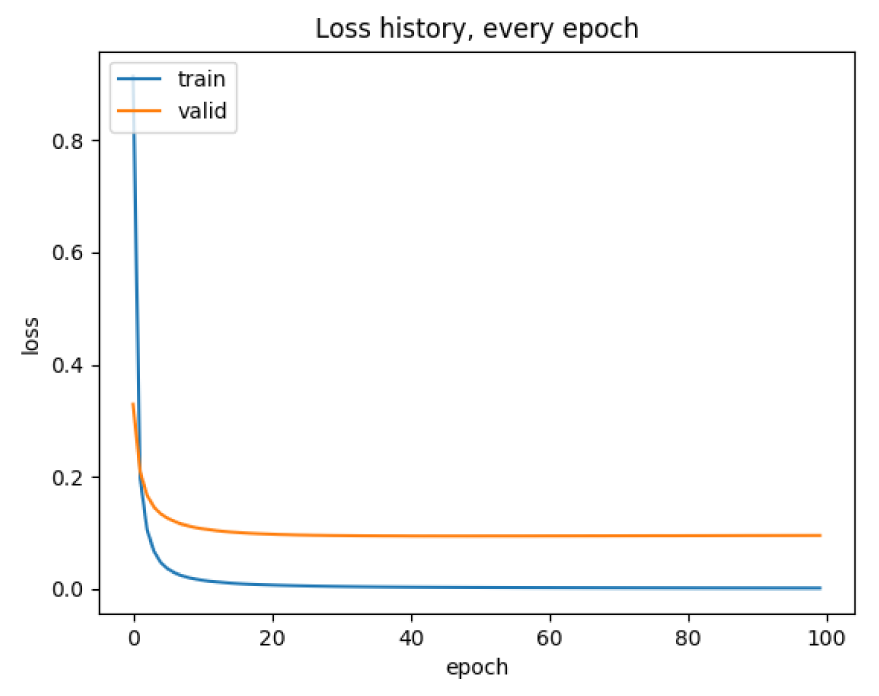
Training and validation loss exhibiting convergence within 100 epochs for the “PT dense 1136 100” model in figure 3.

**Figure S4:**
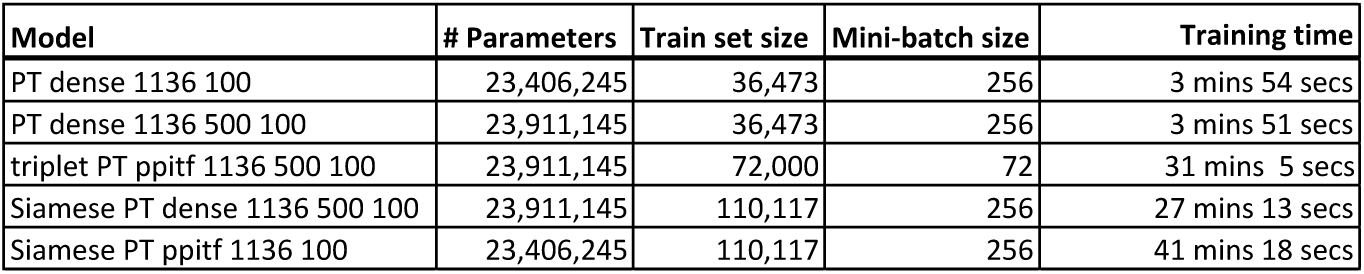
Number of parameters and time to train for each of the neural network model types from Figure 3. All models were trained for 100 epochs on a machine with two Intel(R) Xeon(R) E5-2620 v3 @ 2.40GHz CPUs and a Nvidia GeForce GTX 1080 GPU.

**Figure S5:**
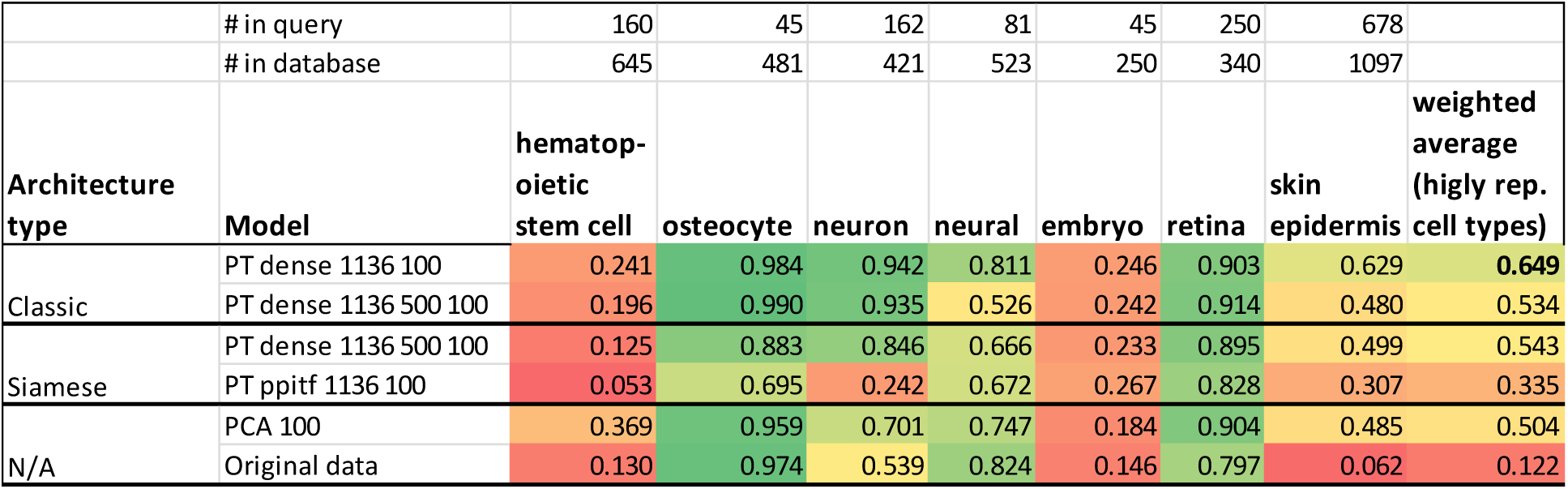
Retrieval testing results of various architectures, as well as PCA and the original (unreduced) expression data, trained on only the data that we processed ourselves. “PT” indicates that the model had been pretrained using the unsupervised strategy (Supporting Methods). Numbers after the model name indicate the hidden layer sizes. For example, “dense 1136 500 100” is an architecture with three hidden layers. The final column is the weighted average score over those cell types with at least 1000 cells in the database (some not shown in table), and the weights are the number of such cells in the query set.

**Figure S6:**
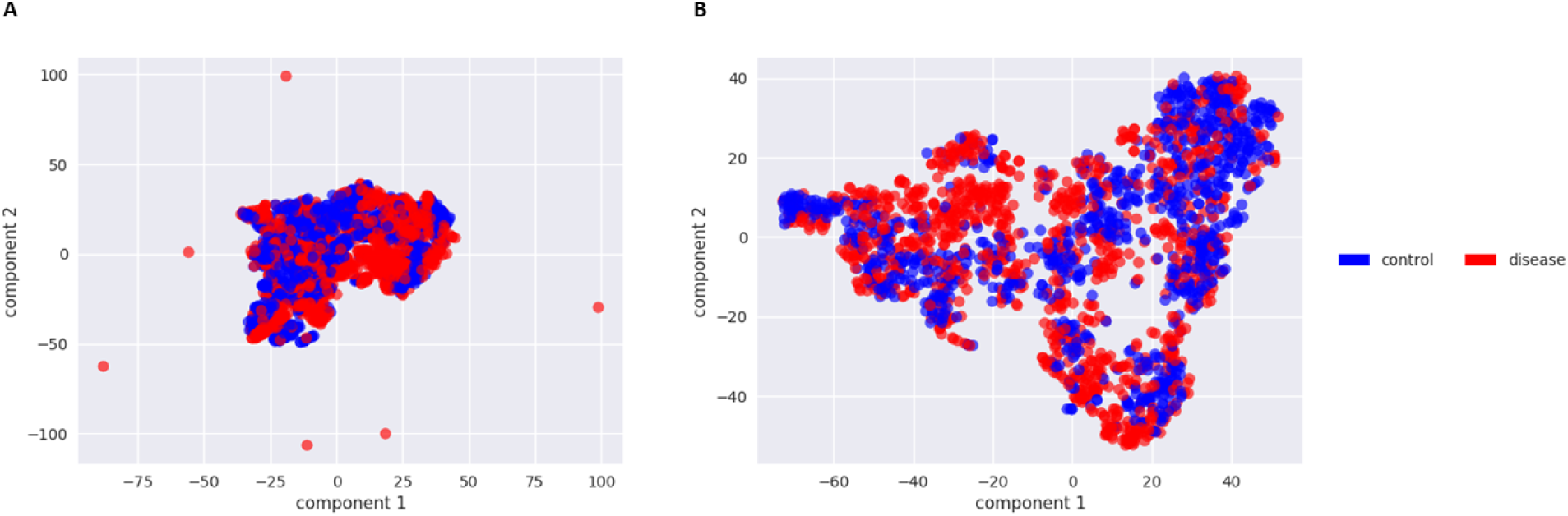
A) Two component t-distributed stochastic neighbor embedding (t-SNE) of the query cells from their original 20,499 dimensions (genes), colored by disease status. B) Two component t-SNE of the query cells from the 100 dimensional neural embedding via the “PT dense 1136 100” model.

**Figure S7:**
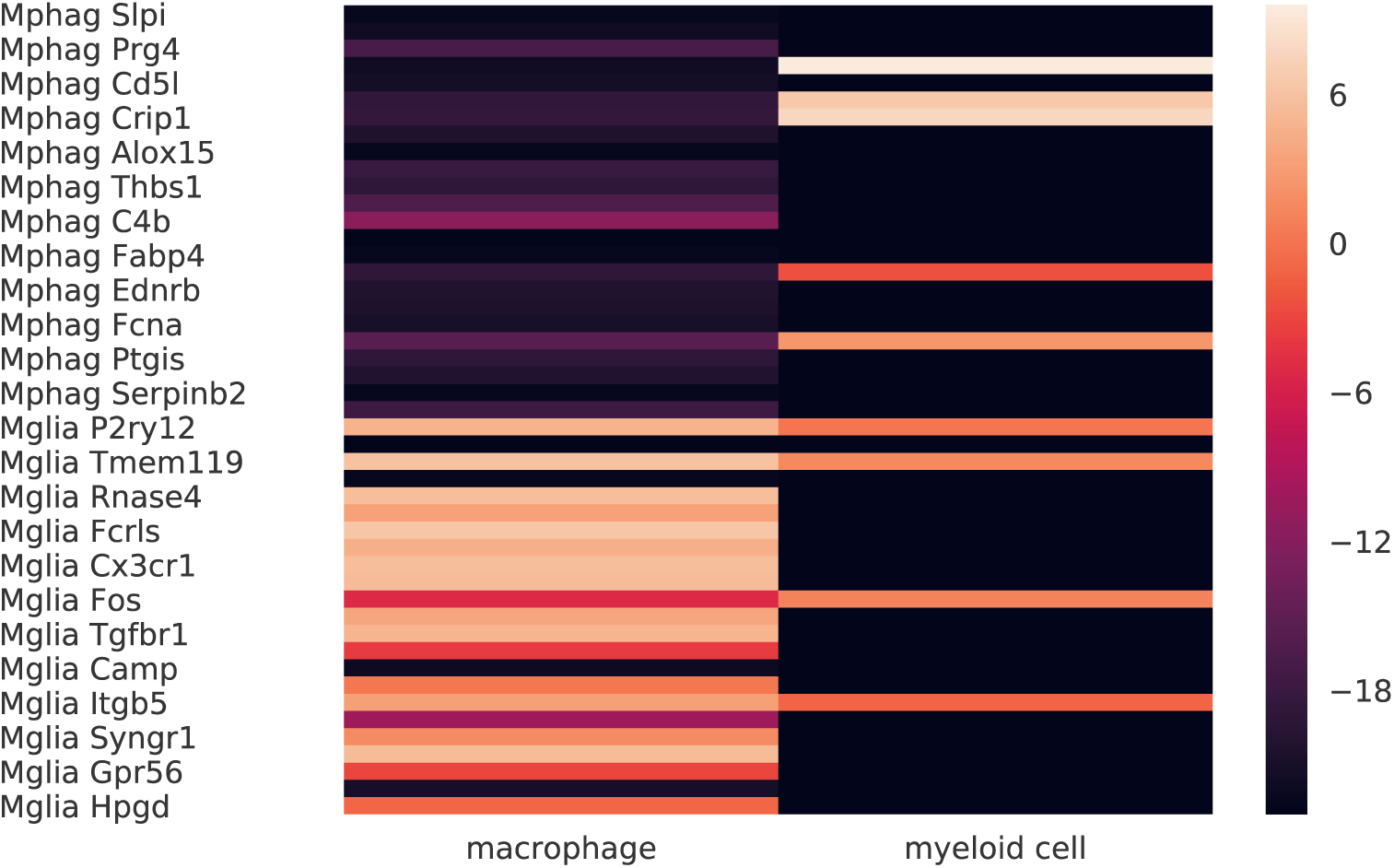
Expression (log_2_(RPKM)) of marker genes (Hickman et al., 2013) for macrophages and microglial cells for all cells we classified as “macrophage” or “myeloid” cell.

**Figure S8:**
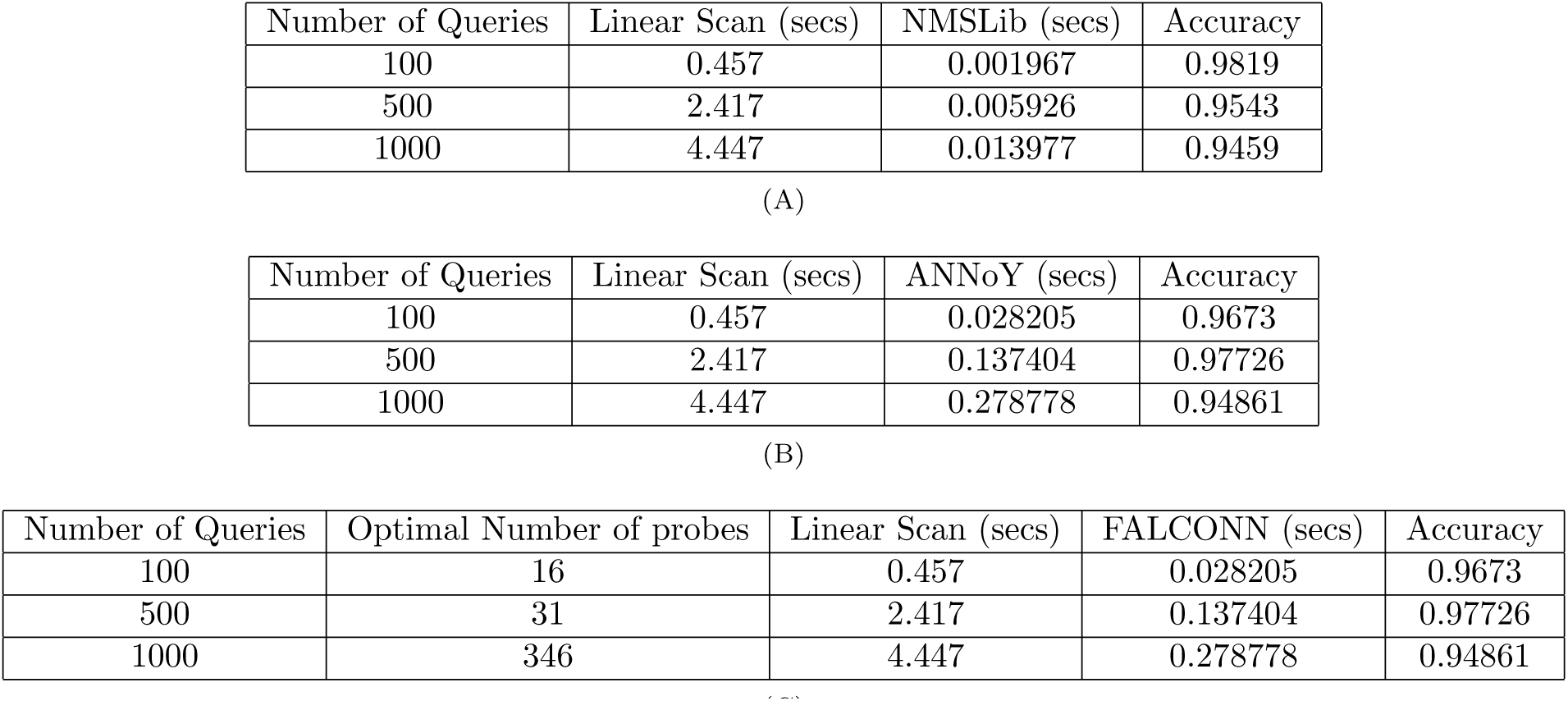
Nearest neighbor search benchmarking results for **(A)** NMSLB, **(B)** ANNoY, and **(C)** FALCONN.

**Algorithm 1:**
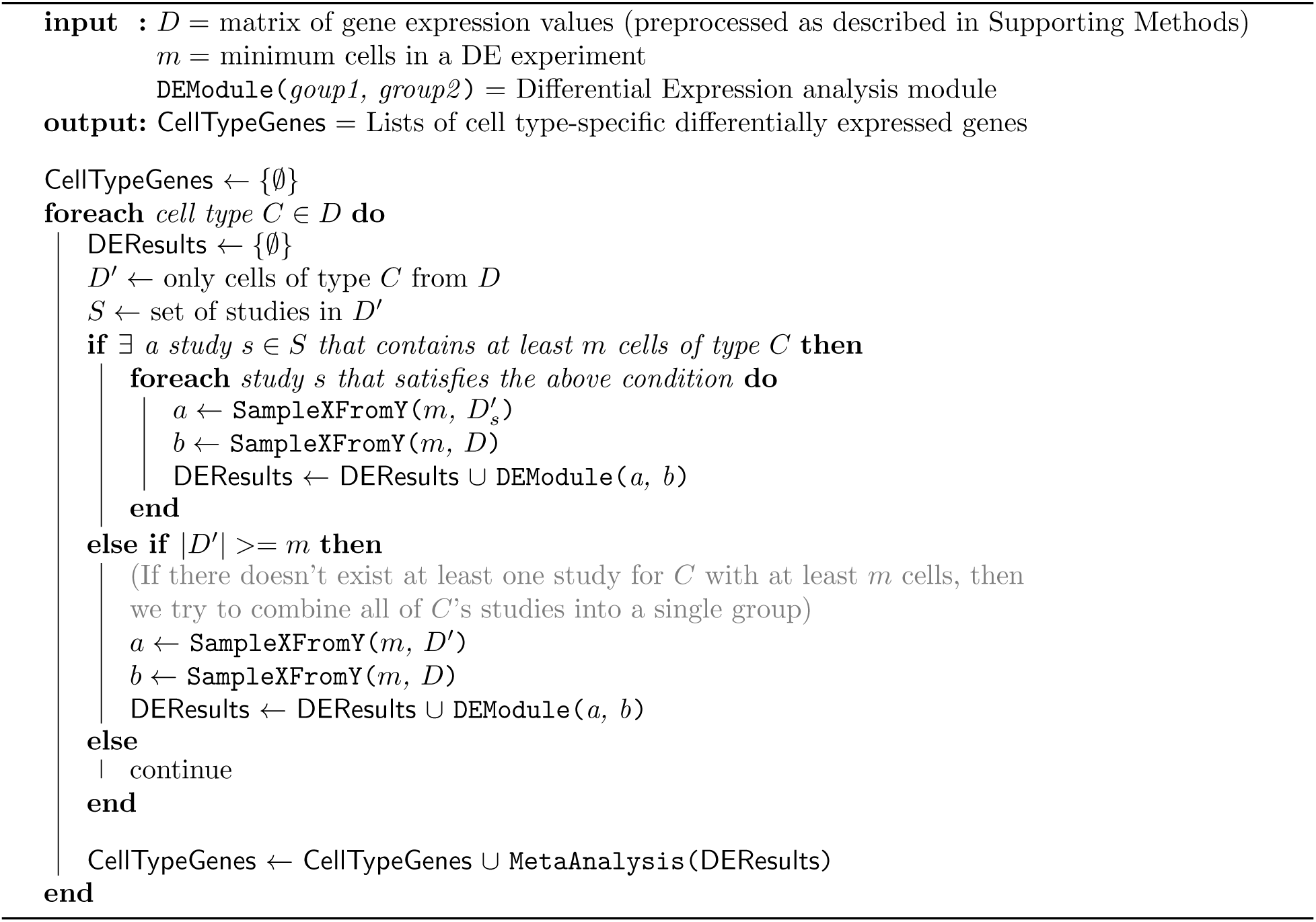
Cell type-specific gene list construction

## Supporting methods

### Triplet architectures trained with batch-hard loss

**Figure S9:**
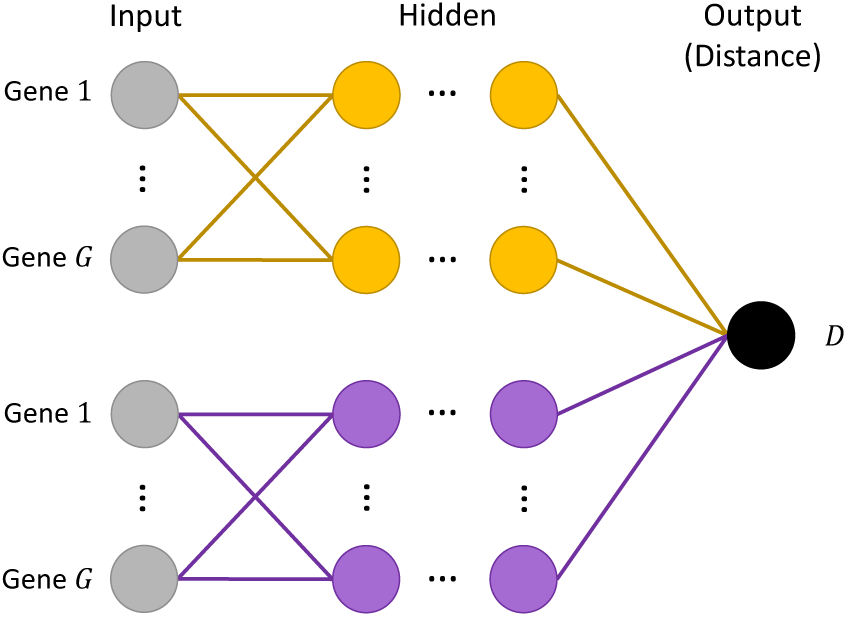
A “Siamese” neural network configuration. Weights are shared between the yellow and the purple subnetworks. Data is fed in pairs with one item in the pair going through the yellow subnetwork, and the other through the purple subnetwork. Triplet networks operate in a similar manner, except that three points are fed through three subnetworks (again, shared weights).

Following the same motivations as Siamese networks, triplet networks also seek to learn an optimal embedding but do so by looking at three samples at a time instead of just two as in a Siamese network. The triplet loss used by Schroff et al. considers an point (anchor), a second point of the same class as the anchor (positive), and a third point of a different class (negative) (Schroff, Kalenichenko, and Philbin, 2015).

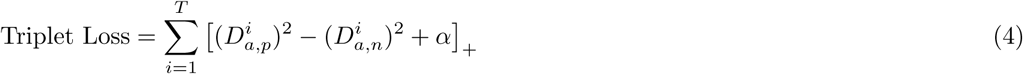

Where:

***T*** is the set of all training examples (triplets of data points)

***D*_*a,p*_**, ***D*_*a,n*_** are the Euclidean distances computed by the network between the embeddings of the anchor point and the positive point, and the embeddings of the anchor point and the negative point, respectively

***α*** is a margin hyperparameter

When training a Siamese or triplet network, it is very important to select pairs/triplets that are difficult enough so that the network learns and converges (Schroff, Kalenichenko, and Philbin, 2015). A straight-forward approach to mine hard examples would involve periodically pausing training, embedding all data points using the current weights, calculating pairwise distances among all of these points in the embedded space, and selecting new examples based on this. This is computationally expensive and is prone to selecting pairs that are too difficult (outliers), so here we have opted to mine challenging pairs in an online fashion by selecting from points within a mini-batch (Hermans, Beyer, and Leibe, 2017; Schroff, Kalenichenko, and Philbin, 2015). Specifically, we use Hermans et al.’s “batch hard” loss, shown in Equation 5 (Hermans, Beyer, and Leibe, 2017). The loss is calculated over a mini-batch, which is constructed by first sampling *P* cell types (labels) and then sampling *K* cells of each type.

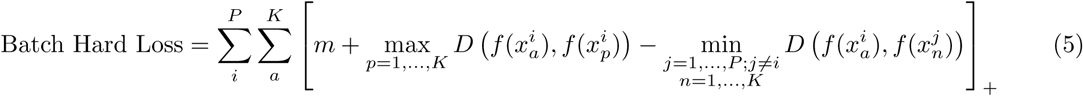

Where:

***f*** is the neural network (an embedding function)

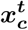is the *c*^*th*^ cell of type *t* in the mini-batch

***D*(*a, b*)** is the Euclidean distance between vectors ***a*** and ***b m*** is a margin hyperparameter

For each anchor point *a* in the mini-batch, the max term in Equation 5 finds the hardest positive pair, while the min term finds the hardest negative pair.

### Unsupervised neural network pretraining

While all of our neural embedding models are trained in a supervised fashion (for cell-type classification or embedding optimization), we also used unsupervised training to initialize the weights for these models, and this was shown to improve performance (Results, Figure 3). We have shown in prior work that using a denoising autoencoder (DAE) for unsupervised pretraining can improve performance (Lin et al., 2017). Here we extend this to using stacks of DAEs, which are each individually trained in a process called “greedy layer-wise pretraining” to pretrain each layer of our neural network architectures (Bengio et al., 2007; Vincent et al., 2010).

Here we illustrate the process with an example of how we would pretrain a neural network with an input layer, two hidden layers, and an output layer (Figure S10a). Figure S10b shows how the first hidden layer is pretrained. The first hidden layer becomes the hidden layer in a DAE, with a “reconstruction” layer of the same dimensionality as the input added on top. Gaussian noise is added to each sample and fed through this DAE, and the training objective is to minimize the mean squared error between the reconstructed output and the clean, uncorrupted samples. After training this DAE, the weights can be used to initialize the first hidden layer in our supervised neural network. This process is repeated for the other hidden layers, where all weights below the layer of interest are fixed, and the input is first fed forward through these fixed weights to get a new representation, and then the corrupting noise is added to this new representation for training the DAE of the current layer.

### Cell-type similarity calculation

Rather than rely on distances based on the ontology graph structure, we computed cell type similarities with a data driven approach by using term co-occurrences in PubMed. For each pair of cell types in our ontology, we queried PubMed for articles which had both of the terms in the title or abstract texts.

We then define *S*_*i*_ as the number of articles in which term *i* occurred in, and *P*_*i,j*_ as the number of articles in which terms *i* and *j* co-occurred in. We additionally define a binary matrix *M* of shape (number of ontology terms, number of documents), where *M*_*i,j*_ = 1 if term *j* occurred in document *i*.

The similarity between two terms can then be computed as a cosine similarity:

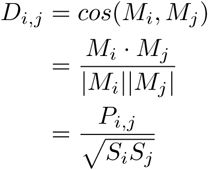

**Figure S10:**
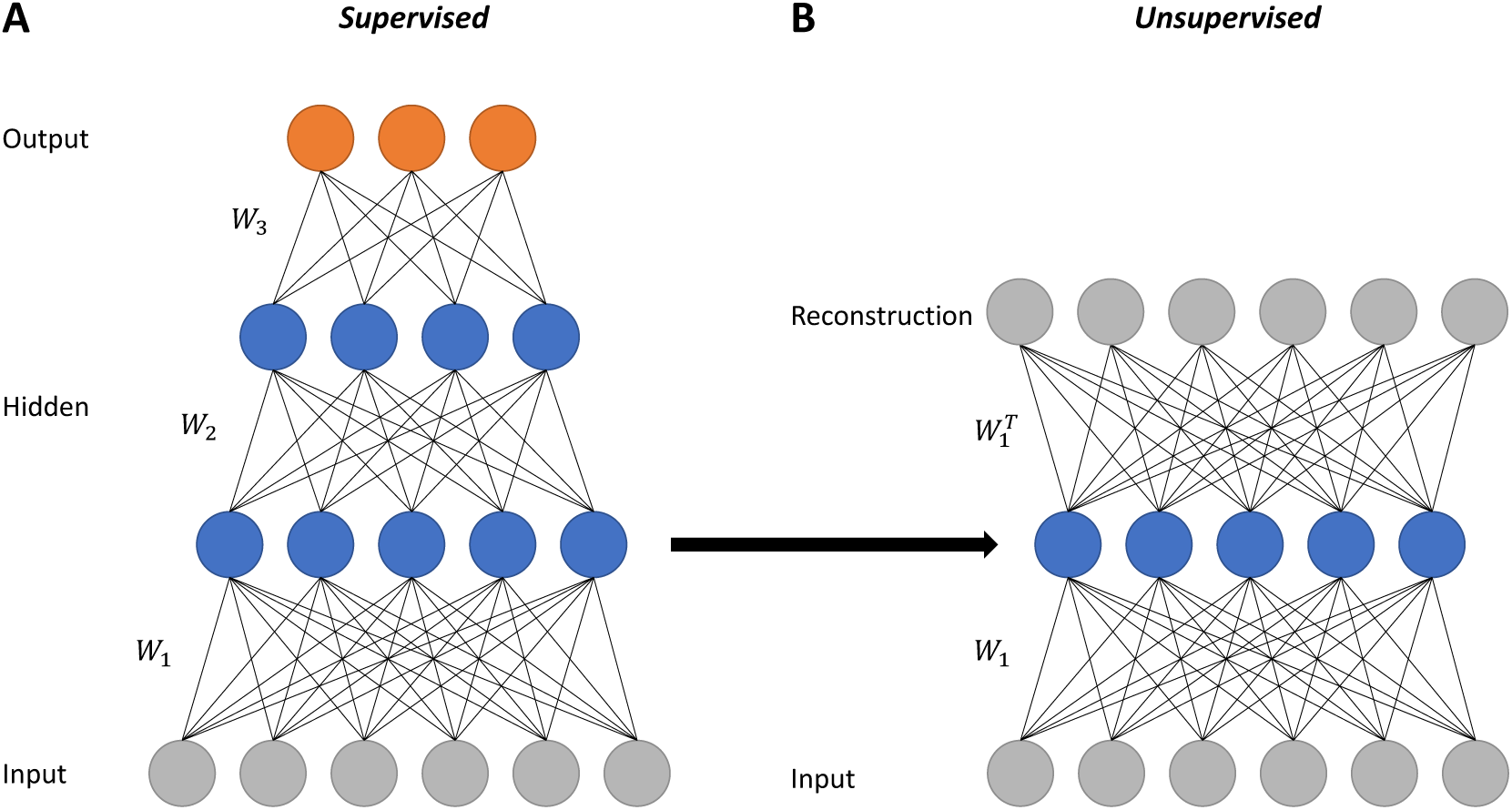
An illustration of one stage of greedy layer-wise unsupervised pretraining of a neural network. *W*_1_, *W*_2_, and *W*_3_ are weight matrices. **(A)** A supervised neural network with two hidden layers to be pretrained. **(B)** An unsupervised DAE that has the the supervised network’s first hidden layer as its middle layer. This DAE is trained to minimize the mean squared error between its reconstruction and the original input data (prior to corruption with noise). After training this DAE, its weights can be used as initial values for the weights in the supervised network.

The above measure is not symmetric, and a symmetric version can be defined as:

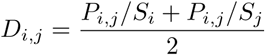

For our neural embedding models that rely on computing distances between cell types (Siamese architectures), we use the symmetric version of the cell type similarities. We use the non-symmetric version to evaluate all of our neural embedding models (*i.e.* to compute the MAFP values in our retrieval analysis).

### Mean average “flexible” precision

We employ a modified version of mean average precision (MAP) called mean average “flexible” precision (MAFP) in order to evaluate the retrieval performance of our dimensionality reduction models. Traditional average precision operates on ranked lists (*e.g.* a list of nearest neighbor cells in a retrieval database) by calculating the precision at exact-match cutoffs in the list, and then taking the mean of these. This requirement is too restrictive for proper evaluation here, as a binary “hit” or “miss” criterion would not allow a list containing similar cell types to be valued higher than lists with less relevant cell types. Instead, average flexible precision is computed by summing the cell-type similarities between the query cell and each retrieved result and dividing by the length of the list at a particular depth, and finally taking the average of these values across all depths in the list.

### Missing data imputation

Our uniform preprocessing pipeline requires the raw reads data for each study. For certain studies, this is not available, and only the RPKM values processed by the authors of the original study are present. We would still like to integrate this data into our database, which requires us to impute the values of the expression of missing genes.

We conduct missing value imputation with a k-nearest neighbors-based approach, similar to procedures in prior work (Lin et al., 2017; Troyanskaya et al., 2001). We ignore cells which have more than 20% of our gene set missing. After filtering those cells out, we impute the remaining values for each study independently. For each study, in order to compute nearest neighbor genes within the study we first fill the missing values with the median expression value in each cell. We then calculate the euclidean distance matrix of the genes in the study (where each gene is represented by a vector of its expression values in each cell). Finally, the value of each missing gene is imputed with the average expression of its 10 nearest neighbor genes within each cell.

### Preprocessing strategies for differential expression analysis

Prior to conducting differential expression analysis, we filter out low-quality genes and cells. To filter out lowly-expressed genes, we first calculate the average reads of each gene within each cell type. We then calculate an overall average for each gene by taking the average of its cell-type specific average expressions. We then only keep genes that have an overall average value that is above a cutoff of 50. This resulted in the removal of 4180 out of 20499 genes.

To filter out cells, we remove cells with fewer than 1.8*×* 103 detected genes, which is the default procedure in the tool we have chosen to use for DE analysis, SCDE (Kharchenko, Silberstein, and Scadden, 2014). This resulted in the removal 732 cells out of 23982 cells. We note that the 23982 cells are a subset of our total database, as we only conduct DE analysis for cell types with a minimum of 75 cells.

### Meta-analysis of differential expression experiments

For cell types that have data from multiple studies, we conduct separate DE analysis for each of those studies, and then combine the results to obtain a single list of differentially expressed genes with their p-values. To do this, we take the maximum FDR adjusted p-value for each gene across the studies as its final p-value. Our list of significant DE genes for a particular cell type is then taken as the set of genes with final p-values below a threshold of 0.05.

Additionally, we only keep the DE genes that had a consistent sign (positive or negative) log_2_ fold-change across the multiple DE analysis that were done for the particular cell type.

### PPI/TF architectures

Since our set of input genes is different, the architectures discussed in Lin et al. could not be used directly here. We used the same transcription factor data as Lin et al. (Schulz et al., 2012). For our protein-protein interaction data, we use a protein-protein interaction network which integrates known and predicted interactions from multiple databases (Szklarczyk et al., 2014).

**Figure S11:**
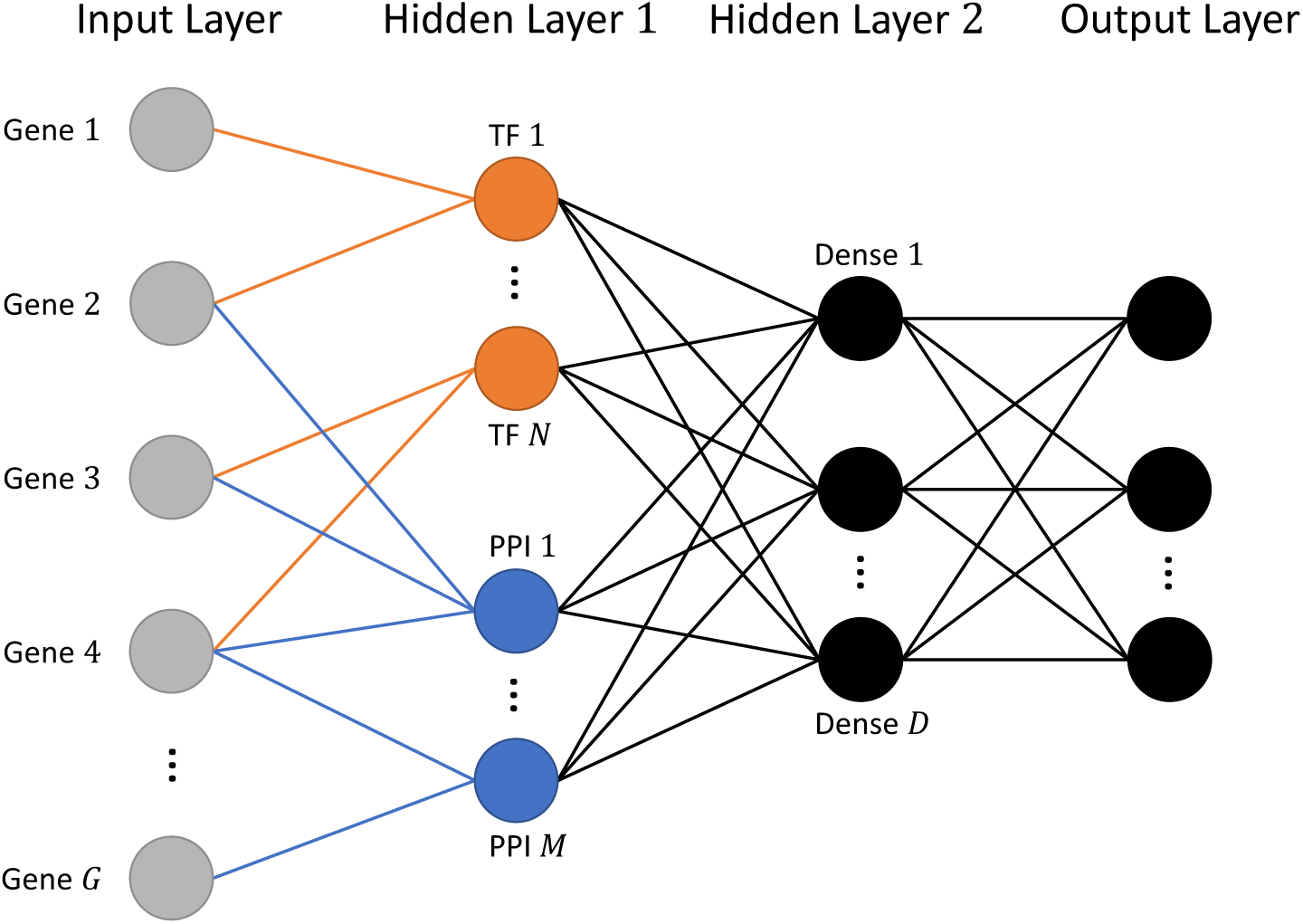
Our Protein-protein/protein-DNA (PPI/TF) based neural network architecture.

### Gene Ontology based architectures

We also tested another method to group genes based on the Gene Ontology (GO) (Consortium, 2017). GO is a directed acyclic graph (DAG) whose nodes are gene products (RNA or protein) and whose edges are relationships between these gene products. Along with this graph, there exist curated annotations which associate genes from specific organisms with each node in the graph. Here we propose that GO’s hierarchical structure will enable our models to recognize complicated relationships that most likely exist between genes.

To construct a hierarchical neural network architecture that mirrors the structure of the GO DAG, we first use the published GO annotations for *M. musculus* (*Mouse Genome Database (MGD)*; Smith et al., 2017) to associate each of our input genes with a node in GO. Specifically, we choose a particular depth *d* (distance from a root node) and first associate the genes with only the nodes at this depth in the GO graph. Multiple genes can be associated with the same node. Then we use this grouping of the input genes as the first hidden layer of a neural network. Each node in this hidden layer represents a GO node at depth d with its associated input genes connected to it. The nodes in the next hidden layer will be constructed from GO nodes that have depth d-1. A node in a layer is connected to a node in the layer above if a path exists between their corresponding nodes in the GO graph. We continue this process until the last hidden layer has the desired number of nodes (the size of our reduced dimension). The final result is the network depicted in Figure S12.

A key point is that the connections in the network are sparse: a node is only connected to a subset of the nodes in the layer below. This is analogous to the idea of convolutional neural networks (CNNs), in which neurons in a given layer are only connected to a subset of neurons in the layer below (the neuron’s “receptive field”) (LeCun et al., 1998).

**Figure S12:**
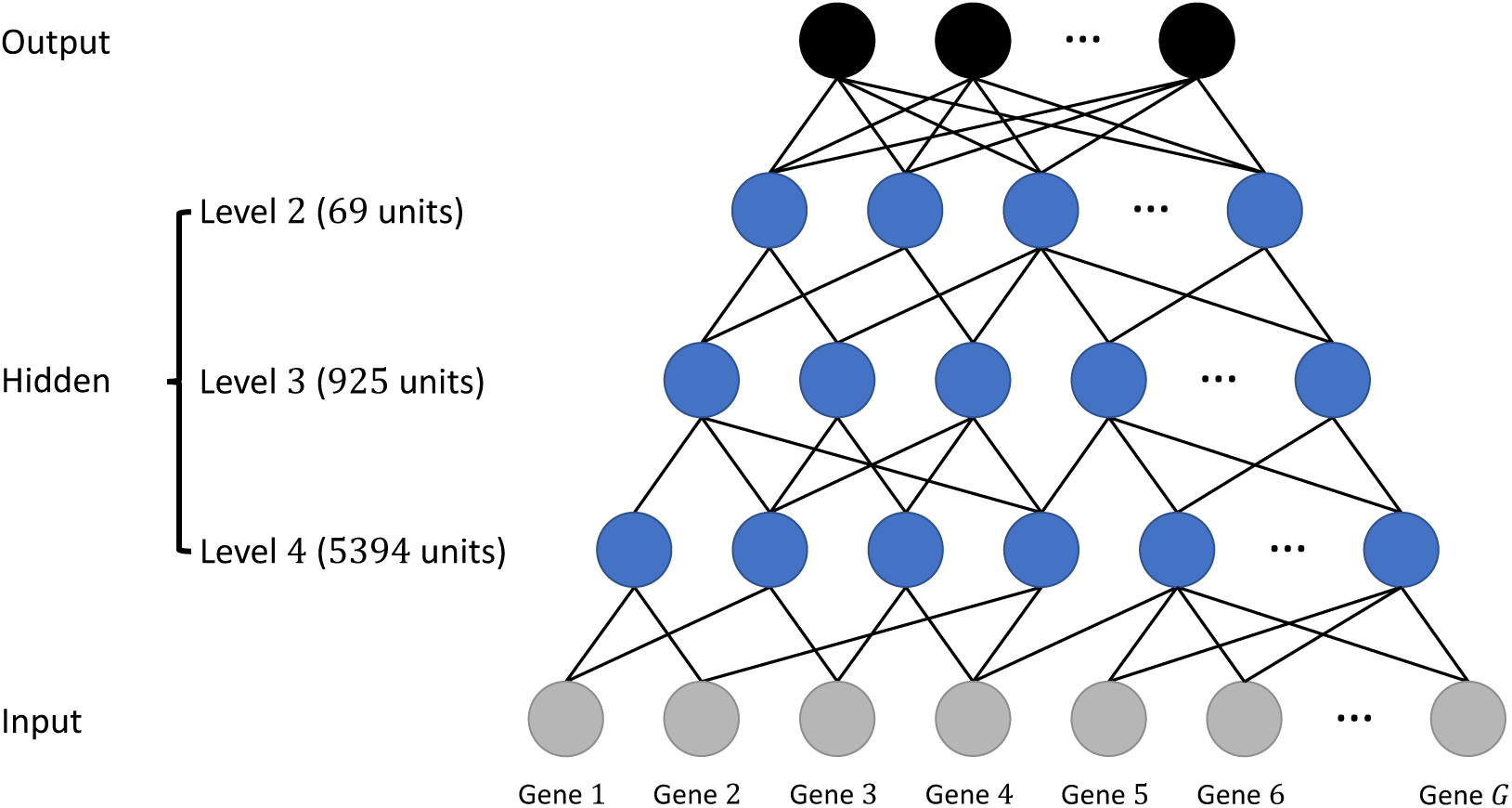
Our Gene Ontology-based neural network architecture. The connections between the input layer and the first hidden layer, as well a the connections between hidden layers, are sparse and follow connections in the GO DAG. The final hidden layer is fully connected (dense) to the output layer.

## Footnotes

1 https://github.com/spotify/annoy

## References

Andoni, Alexandr et al. (2015). “Practical and optimal LSH for angular distance”. In: Advances in Neural Information Processing Systems, pp. 1225–1233.

Bard, Jonathan, Seung Y Rhee, and Michael Ashburner (2005). “An ontology for cell types”. In: Genome biology 6.2, R21.

Bengio, Yoshua et al. (2007). “Greedy layer-wise training of deep networks”. In: Advances in neural information processing systems, pp. 153–160.

Boytsov, Leonid and Bilegsaikhan Naidan (2013). “Engineering Efficient and Effective Non-metric Space Library”. In: Similarity Search and Applications - 6th International Conference, SISAP 2013, A Coruña, Spain, October 2-4, 2013, Proceedings, pp. 280–293. DOI:10.1007/978-3-642-41062-8_28. url: https://doi.org/10.1007/978-3-642-41062-8_28.

Bullard, James H et al. (2010). “Evaluation of statistical methods for normalization and differential expression in mRNA-Seq experiments”. In: BMC bioinformatics 11.1, p. 94.

Chollet, François et al. (2015). Keras. https://github.com/fchollet/keras.

Chopra, Sumit, Raia Hadsell, and Yann LeCun (2005). “Learning a similarity metric discriminatively, with application to face verification”. In: Computer Vision and Pattern Recognition, 2005. CVPR 2005. IEEE Computer Society Conference on. Vol. 1. IEEE, pp. 539–546.

Consortium, Gene Ontology et al. (2017). “Expansion of the Gene Ontology knowledgebase and resources”. In: Nucleic acids research 45.D1, pp. D331–D338.

Hermans, Alexander, Lucas Beyer, and Bastian Leibe (2017). “In defense of the triplet loss for person reidentification”. In: arXiv preprint arXiv:1703.07737.

Hickman, Suzanne E et al. (2013). “The microglial sensome revealed by direct RNA sequencing”. In: Nature neuroscience 16.12, p. 1896.

Hinton, Geoffrey E and Ruslan R Salakhutdinov (2006). “Reducing the dimensionality of data with neural networks”. In: science 313.5786, pp. 504–507.

Jaitin, Diego Adhemar et al. (2014). “Massively parallel single-cell RNA-seq for marker-free decomposition of tissues into cell types”. In: Science 343.6172, pp. 776–779.

Kharchenko, Peter V, Lev Silberstein, and David T Scadden (2014). “Bayesian approach to single-cell differential expression analysis”. In: Nature methods 11.7, p. 740.

Kim, D., B. Langmead, and S. L. Salzberg (2015). “HISAT: a fast spliced aligner with low memory requirements”. In: Nat. Methods 12.4, pp. 357–360.

Koch, Gregory, Richard Zemel, and Ruslan Salakhutdinov (2015). “Siamese neural networks for one-shot image recognition”. In: ICML Deep Learning Workshop. Vol. 2.

Kolodziejczyk, Aleksandra A. et al. (2015). “The Technology and Biology of Single-Cell RNA Sequencing”. In: Molecular Cell 58.4, pp. 610–620. issn: 1097-2765. DOI:https://doi.org/10.1016/j.molcel.2015.04.005. url: http://www.sciencedirect.com/science/article/pii/S1097276515002610.

LeCun, Yann et al. (1998). “Gradient-based learning applied to document recognition”. In: Proceedings of the IEEE 86.11, pp. 2278–2324.

Lescroart, Fabienne et al. (2018). “Defining the earliest step of cardiovascular lineage segregation by singlecell RNA-seq”. In: Science, eaao4174.

Lin, Chieh et al. (2017). “Using neural networks for reducing the dimensions of single-cell RNA-Seq data”. In: Nucleic Acids Research 45.17, e156. DOI:10.1093/nar/gkx681. eprint: oup/backfile/content_public/journal/nar/45/17/10.1093_nar_gkx681/2/gkx681.pdf. url: +http://dx.doi.org/10.1093/nar/gkx681.

Mathys, Hansruedi et al. (2017). “Temporal Tracking of Microglia Activation in Neurodegeneration at SingleCell Resolution”. In: Cell reports 21.2, pp. 366–380.

Mouse Genome Informatics, The Jackson Laboratory. Mouse Genome Database (MGD). url: http://www.informatics.jax.org (visited on 07/10/2017).

Park, Thomas In-Hyeup et al. (2012). “Adult human brain neural progenitor cells (NPCs) and fibroblast-like cells have similar properties in vitro but only NPCs differentiate into neurons”. In: PloS one 7.6, e37742.

Patel, Anoop P et al. (2014). “Single-cell RNA-seq highlights intratumoral heterogeneity in primary glioblastoma”. In: Science, p. 1254257.

Pierson, Emma and Christopher Yau (2015). “ZIFA: Dimensionality reduction for zero-inflated single-cell gene expression analysis”. In: Genome biology 16.1, p. 241.

Rizvi, Abbas H et al. (2017). “Single-cell topological RNA-seq analysis reveals insights into cellular differentiation and development”. In: Nature biotechnology 35.6, p. 551.

Robinson, Mark D and Alicia Oshlack (2010). “A scaling normalization method for differential expression analysis of RNA-seq data”. In: Genome biology 11.3, R25.

Rosenbloom, Kate R et al. (2014). “The UCSC genome browser database: 2015 update”. In: Nucleic acids research 43.D1, pp. D670–D681.

Schroff, Florian, Dmitry Kalenichenko, and James Philbin (2015). “Facenet: A unified embedding for face recognition and clustering”. In: Proceedings of the IEEE conference on computer vision and pattern recognition, pp. 815–823.

Schulz, Marcel H et al. (2012). “DREM 2.0: Improved reconstruction of dynamic regulatory networks from time-series expression data”. In: BMC systems biology 6.1, p. 104.

Smith, Cynthia L et al. (2017). “Mouse Genome Database (MGD)-2018: knowledgebase for the laboratory mouse”. In: Nucleic acids research 46.D1, pp. D836–D842.

Szklarczyk, Damian et al. (2014). “STRING v10: protein–protein interaction networks, integrated over the tree of life”. In: Nucleic acids research 43.D1, pp. D447–D452.

Tirosh, Itay et al. (2016). “Dissecting the multicellular ecosystem of metastatic melanoma by single-cell RNA-seq”. In: Science 352.6282, pp. 189–196.

Troyanskaya, Olga et al. (2001). “Missing value estimation methods for DNA microarrays”. In: Bioinformatics 17.6, pp. 520–525.

Usoskin, Dmitry et al. (2015). “Unbiased classification of sensory neuron types by large-scale single-cell RNA sequencing”. In: Nature neuroscience 18.1, p. 145.

Vanlandewijck, Michael et al. (2018). “A molecular atlas of cell types and zonation in the brain vasculature”. In: Nature 554.7693, p. 475.

Vincent, Pascal et al. (2010). “Stacked denoising autoencoders: Learning useful representations in a deep network with a local denoising criterion”. In: Journal of Machine Learning Research 11.Dec, pp. 3371–3408.

Wills, Quin F et al. (2013). “Single-cell gene expression analysis reveals genetic associations masked in whole-tissue experiments”. In: Nature biotechnology 31.8, pp. 748–752.

Yau, Christopher et al. (2016). “pcaReduce: hierarchical clustering of single cell transcriptional profiles”. In: BMC bioinformatics 17.1, p. 140.

Zeisel, Amit et al. (2015). “Cell types in the mouse cortex and hippocampus revealed by single-cell RNA-seq”. In: Science 347.6226, pp. 1138–1142.

Zhang, Ye et al. (2014). “An RNA-sequencing transcriptome and splicing database of glia, neurons, and vascular cells of the cerebral cortex”. In: Journal of Neuroscience 34.36, pp. 11929–11947.

